# A MALDI-MS biotyping-like method to address honey bee health status through computational modelling

**DOI:** 10.1101/683607

**Authors:** Karim Arafah, Sébastien Nicolas Voisin, Victor Masson, Cédric Alaux, Yves Le Conte, Michel Bocquet, Philippe Bulet

## Abstract

Among pollinator insects, bees undoubtedly account for the most important species. They play a critical role in boosting reproduction of wild and commercial plants and therefore contribute to the maintenance of plant biodiversity and sustainability of food webs. In the last few decades, domesticated and wild bees have been subjected to biotic and abiotic threats, alone or in combination, causing various health disorders. Therefore, monitoring solutions to improve bee health are increasingly necessary. MALDI mass spectrometry has emerged within this decade as a powerful technology to biotype micro-organisms. This method is currently and routinely used in clinical diagnosis where molecular mass fingerprints corresponding to major protein signatures are matched against databases for real-time identification. Based on this strategy, we developed MALDI BeeTyping as a proof of concept to monitor significant hemolymph molecular changes in honey bees upon infection with a series of entomopathogenic Gram-positive and -negative bacteria. A *Serratia marcescens* strain isolated from one “naturally” infected honey bee collected from the field was also considered. We performed a series of individually recorded hemolymph molecular mass fingerprints and built, to our knowledge, the first computational model made of nine molecular signatures with a predictive score of 97.92%. Hence, we challenged our model by classifying a training set of individual bees’ hemolymph and obtained overall recognition of 91.93%. Through this work, we aimed at introducing a novel, realistic, and time-saving high-throughput biotyping-like strategy that addresses honey bee health in infectious conditions and on an individual scale through direct “blood tests”.

**Significance Statement:** Domesticated and wild bees worldwide represent the most active and valuable pollinators that ensure plant biodiversity and the success of many crops. These pollinators and others are exposed to deleterious pathogens and environmental stressors. Despite efforts to better understand how these threats affect honey bee health status, solutions are still crucially needed to help beekeepers, scientists and stakeholders in obtaining either a prognosis, an early diagnosis or a diagnosis of the health status of the apiaries. In this study, we describe a new method to investigate honey bee health by a simple “blood test” using fingerprints of some peptides/proteins as health status signatures. By computer modelling, we automated the identification of infected bees with a predictive score of 97.92%.

## Introduction

Over several decades, an abnormal mortality of honey bees and other pollinators (bumblebees, solitary bees) has been observed in all industrialized countries (1–4). This phenomenon has been particularly recorded in honey bees (5). The global loss of honey bee colonies has detrimental consequences for plant biodiversity, bee products, and negative economic and societal effects (6). As a result, many scientific studies have been carried out to understand the mechanisms underlying phenomena such as colony weakening or collapse and colony mortality observed in most of the countries practicing intensive agriculture. Many reports concluded that biotic and abiotic factors are suspected to be involved in this phenomenon, either alone or in combination (2, 5, 7–10). Potential causes are exposure to (i) environmental and in-hive chemicals (11, 12), (ii) agricultural practices (13, 14), (iii) infection by micro-organisms and predation by parasites (15–17) and (iv) nutritional factors (18–20), among others, which lead to the transition from a health status qualified as normal to a health decline that would contribute to the colony collapse (7). The expression of this pathological state may notably be linked to a decrease in the immune capacities of the bee and/or the colony subjected to these combinations of stressors (21–25). The complex underlying mechanisms of stressors (biotic and abiotic) that affect bees and impact honey bee health status remain still partially understood. Both the fundamental molecular mechanisms associated with the modifications of health status and the development of solutions capable of rendering a prognosis, an early diagnosis or a diagnosis, remain to be elucidated. This is a prerequisite for limiting colony losses and protecting honey bees, but the tools and services to perform a clear sanitary diagnosis of beehives are currently lacking. Even if visual and PCR analyses are available for surveillance of pathogen loads, prediction of the likely impact on the colony remains an issue not satisfactorily addressed (26). Apart from typical methods for honey bee colony health monitoring like polymerase chain reaction assays and sensor-based devices (27–29), mass spectrometry (MS), which has been greatly improved in the past 20 years, may play an essential role in the quest for innovative solutions in monitoring bee health. Among the different MS approaches, Matrix-Assisted Laser Desorption-Ionization – Time of Flight Mass Spectrometry (MALDI-TOF MS) has become increasingly popular for biological sample identification in laboratory research and for clinical diagnostics in microbiology. The widespread interest of this technology for analysing biological matrices is due to the generation of mostly monocharged ions which satisfy the generation of simplified mass spectra when analysing complex biological samples (30).

In the past ten years, MALDI-TOF MS has become a referenced system in microbiological laboratories. Technological developments, making this analytical technique a robust, fast and widely used commercial platform, paved the way for its use in routine clinical microbiology (31–33). In 2013, two independent systems, the VITEK^®^ mass spectrometer (bioMérieux clinical Diagnostics) and the MALDI Biotyper® MicroFlex (Bruker Daltonics Inc.) received the US Food and Drug Administration (FDA) clearance for the identification by biotyping of micro-organism species including yeasts and aerobic / anaerobic bacteria. In 2015, FDA clearance was announced for 193 and over 280 species using the VITEK^®^ and the MALDI Biotyper^®^ instruments, respectively (34, 35). These platforms perform by targeting the ribosomal proteins as biomarkers for the identification of clinical bacterial, fungal and yeast isolates (36). In early 2018, the Bruker MALDI Biotyper^®^ solution received international approvals as an official method of analysis for the food industry (source from Bruker Corporation, related link https://www.bruker.com). Barcoding or molecular mass fingerprints (MFP) of biological matrices by MALDI-TOF MS is indeed a thriving approach, enabling the rapid detection of peptide/protein components that can provide comparative information.

Building on the concept of this MS-based MFP approach and on the demonstrated capability of MALDI-TOF MS to decipher the molecular mechanisms of insect immunity in the *Drosophila* model for various infections (37–39), we performed a peptidomics/proteomics-based mass fingerprinting of honey bee hemolymph using MALDI MS to discriminate different models of bacterial infections. Relying on previously published studies (39–42), we first developed and validated an experimental model of challenged honey bees with Gram-positive and -negative bacteria (using notably a *Serratia marcescens* strain isolated from honey bees). Then, we assessed the usefulness of MALDI-TOF MS to fingerprint the peptides/proteins in honey bee hemolymph in order to build and validate a computational model of bacterial recognition based on the molecular signatures within the molecular mass range 2-20 kDa; a method we will refer to as BeeTyping. In addition, we determined the performance of this computational model using a training set of challenged bees. Through this work, we introduce BeeTyping as an effective method for monitoring honey bee health status by diagnosing bacterial infections in young adult honey bees.

## Results & Discussion

### MALDI-TOF MS Biotyping successfully diagnosed a bacterial strain isolated from honey bee hemolymph

In order to generate relevant biological models of honey bee infections, we used the bacteria *Micrococcus luteus*, *Pectobacterium carotovorum* subsp*. carotovorum* and *Serratia entomophila. M. luteus* is a Gram-positive bacterial strain frequently used when monitoring insect immunity (43) and has been shown to colonize bee hives and gastrointestinal tracts of honey bees (44) while *P. carotovorum* subsp. *carotovorum* (45) and *S. entomophila* (Institut Pasteur, CIP102919) are two bacterial strains that trigger a systemic immune response in insects. We also performed an additional model of infection using a *Serratia marcescens* strain (*Sm*BIOP160412, Lab. collection) isolated from a naturally infected honey bee collected in the field. To certify the constructed biological models of infection, the four bacterial strains were classified by MALDI MS biotyping (Figure 1). As shown, the individual MFPs of the different strains detailed below and represented by the spectral gel views passed the threshold score of identification with a significant score (reliability score ≥ 2) and successfully matched the Bruker reference strains (see Fig. 1, *M. luteus* ref. 1270, *P. carotovorum* subsp. *carotovorum* ref. 78398, and *S. entomophila* ref. 42906) of the MALDI BioTyper® database. Moreover, applied to the bacterial strain *Sm*BIOP160412 isolated and cultured from an isolate obtained from a naturally infected honey bee, MALDI MS biotyping demonstrated for the first time to our knowledge, its ability to characterize a field bacterial infection in *Apis mellifera*. This identified bacterial strain, namely *S. marcescens*, is known to be a widespread pathogen of adult honey bees (46), and a virulent opportunist, taking advantage of disturbed microbiota to develop in honey bee guts after exposure to the pesticide glyphosate (47).

**Figure 1:**
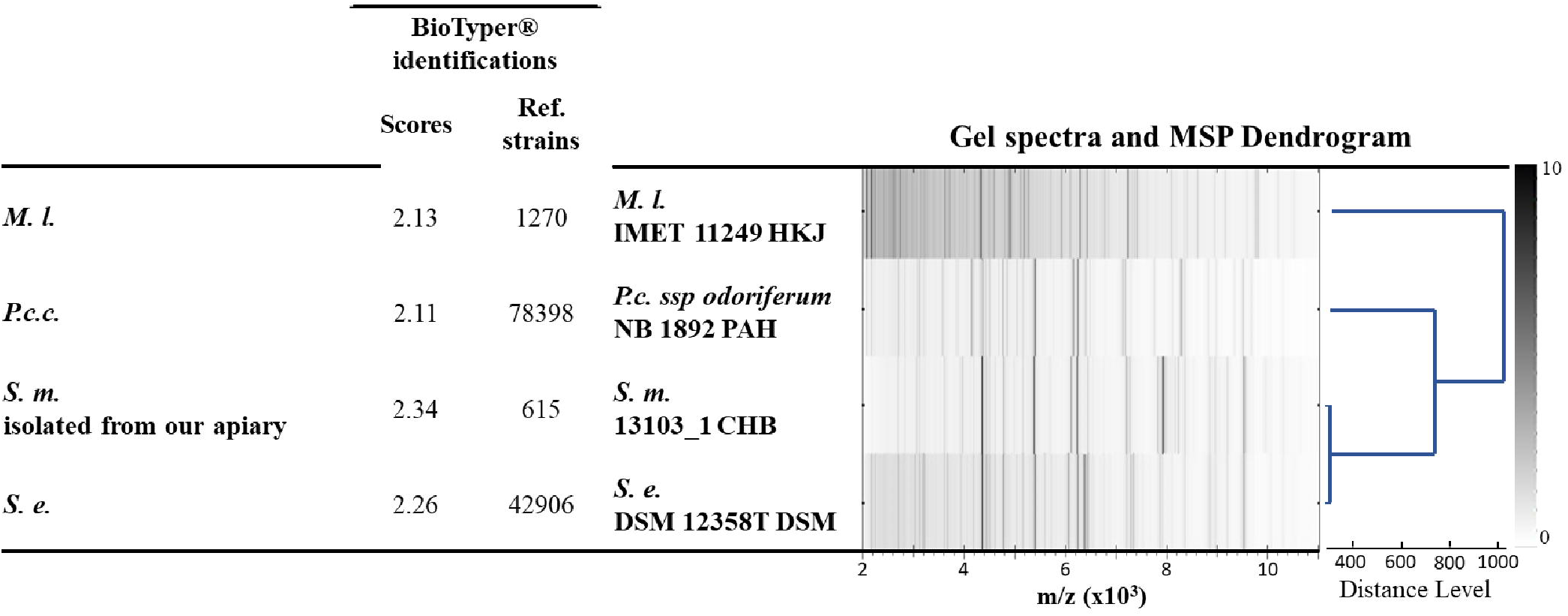
Classification by MALDI-MS biotyping of *Micrococcus luteus* (*M. l.*) and *Pectobacterium caratovorum* subsp. *caratovorum, (P. c. c.)*, *Serratia entomophila*, (*S. e.*) bacteria including the *Serratia marcescens* (*S. m*.) isolated from a naturally infected bee. The strains used to build the infectious model (*M. l.* and *P. c.c.*) and to assess the computational capability to discriminate proteomic fingerprints from the same bacterial species (*S. m.* and *S. e.*). were analyzed in order to confirm their identity based on their molecular profiles by matching the MALDI MS spectra (mass range *m*/*z* 2,000 to 11,000) to the reference strains of the Bruker database containing 6,903 MSP. The BioTyper parameters to validate the identifications of strains, *i.e.* the scores and the matching strains (references and library), were obtained for each biotyped bacterial sample in addition to the MALDI MS spectral gel-view. The dendrogram was built from the Main Spectra Projection (MSP) statistical mode of calculation, which is used to identify, analyze and classify the MALDI MS spectra. These bacteria were identified by their scores and classified according to their distance level (MSP Dendogram) in comparison to reference strains from the database MBT Compass 4.1, build 70 (*M. l*. IMET 11249HKJ, *P. c. ssp odoriferum* NB 1892 PAH, *S. m.* 13103_1 CHB and *S. e.* DSM 12358T DSM). The mass spectra *(m*/*z*) were transformed into gel views where the grey scale bar and thickness of the lines refer to the *m*/*z* peak intensities. Classically, a high confidence identification is obtained with a score between 2.00 and 3.00, a low confidence identification with a score between 1.70 and 1.99 and a failed identification with a score strictly below 1.70. The mass spectra (m/z) were transformed into gel views where the grey scale bar and thickness of the lines refer to the m/z peak intensities.

### MALDI-TOF MS BeeTyping as a new approach to discriminate Gram-positive and -negative biological models of infection directly from honey bee hemolymph

A set of 64 MALDI MS spectra was recorded from individual hemolymph samples. These spectra were obtained from 22 control honey bees and from 23 and 19 honey bees individually infected with the Gram-positive *M. luteus* or the Gram-negative *P. carotovorum* subsp. *carotovorum*, respectively. An averaged spectrum, containing 110 MALDI MS ion peaks (MFP, Table S1), was built for each of the three biological models (Figure 2A). Statistical analysis based on Principal Component Analysis (PCA) and performed on these MFPs clearly segregated the three biological models (Figure 2B). As shown by the PCA plot score, the individual spectra were clustered in accordance with their corresponding models and were segregated based on their mass fingerprints. The unsupervised hierarchical clustering of hemolymph samples, classified almost all of the individual MFPs with respect to their corresponding biological models (Figure 2C). Out of 64 normalized spectra used to build the clustering dendrogram, four and three recorded mismatched spectra were observed, corresponding to the lowest (70%) and highest (95%) limits of explained variance, respectively. The mismatched spectra were further identified within the representation of the PCA plot score of the hemolymph samples (Figure 2B, arrows). At the limit of explained variance of 70%, the four spectra included one spectrum from the control condition and one from the *M. luteus* infection model, both classified under the *P. carotovorum* subsp. *carotovorum* model, and two of this latter model, mismatched to the control model (see asterisks in Fig. 2C). Regarding the three mismatched spectra observed at 95% of the explained variance, one was from the *M. luteus* model and classified under the *P. carotovorum* subsp. *carotovorum* model and two, from the *P. carotovorum* subsp. *carotovorum* model, classified under the control model (see asterisks in Fig. 2C).

**Figure 2:**
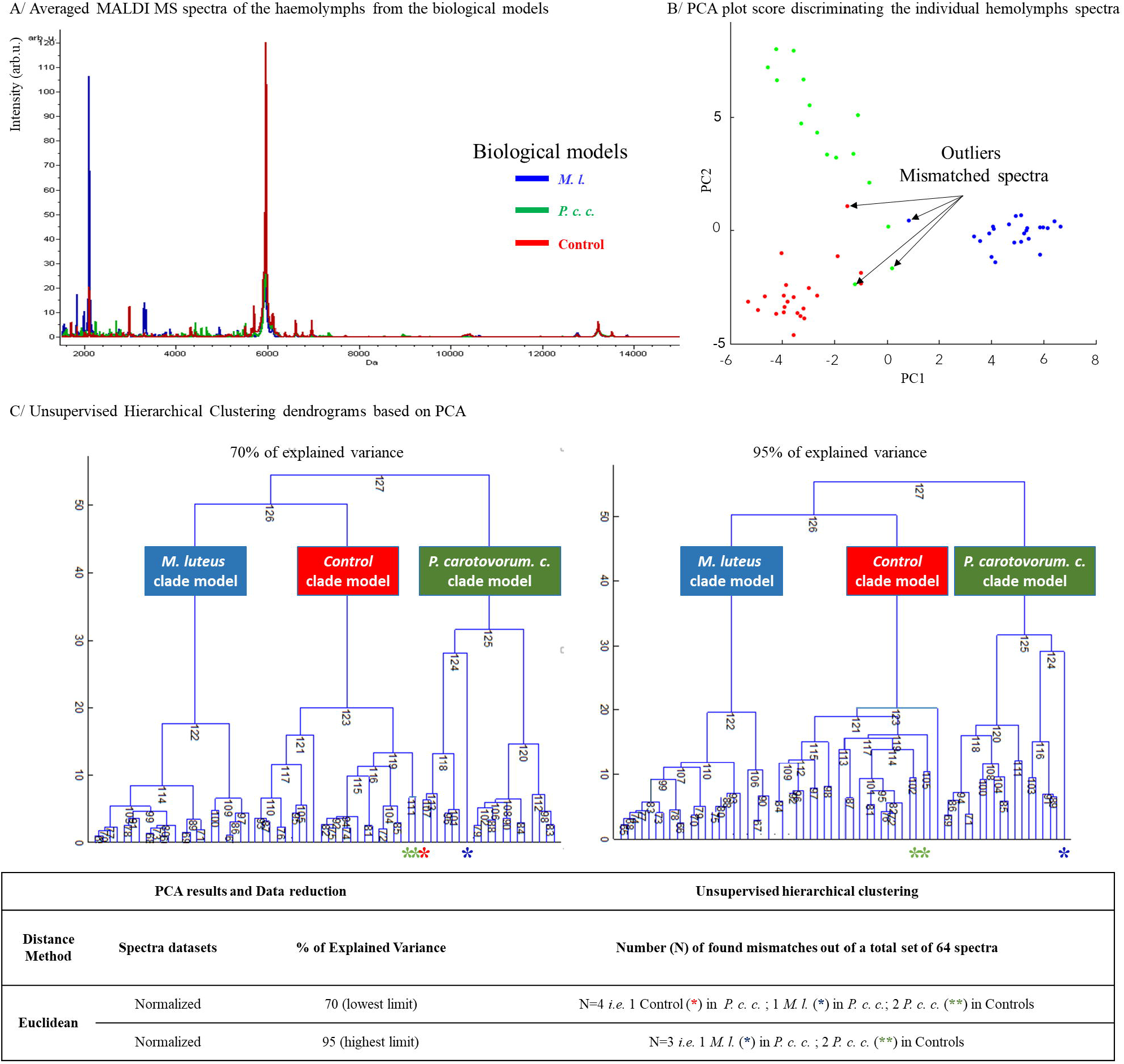
Differential PCA-based statistical analyses and hierarchical clustering of individual hemolymph samples from the biological models. Total averaged spectra were fingerprinted by MALDI MS from the infected and control individuals (**A**). The individual spectra were subjected to PCA analysis, which discriminated the hemolymph molecular mass fingerprints of *Micrococcus luteus* (*M. l.* in blue), *Pectobacterium caratovorum* subsp. *carotovorum* 15 (*P. c. c.* in green) and control (red) groups (**B**); arrows mark the mismatched outliers. An unsupervised hierarchical clustering based on the PCA results classified the individual spectra according to the lowest (70% left panel **C**) and highest (95%, right panel **C**) limits of explained variances.

In order to assess the relationship between the MALDI-TOF MS MFPs of the biological models and the honey bee’s immune status, we correlated these MFPs with each of the four antimicrobial peptides (AMP) defined from *Apis mellifera* (48): Apidaecin 1A (41) at *m*/*z* 2,107), Hymenoptaecin (49) at *m*/*z* 10,270, Abaecin (50) at *m*/*z* 3,878 and Defensin 1A (40) at *m*/*z* 5,519 (Figure 3). As shown, per-peak fingerprint correlations with the antimicrobial peptides (AMPs) were scored based on the molecular ion peak area and represented as heat maps through a colored scale intensity ranging from low (minimum score of −1, in red) to high correlation (maximum score of 1, in green). Reported in relation to the MFPs, four clades (A, B, C and D) described the positive and negative correlations of the MFPs with each of the four AMPs (see Table S1) and segregated the three biological models (non-experimentally infected as a control condition, *P. carotovorum* subsp. *carotovorum.* model and *M. luteus* model, see Fig. 3).

**Figure 3:**
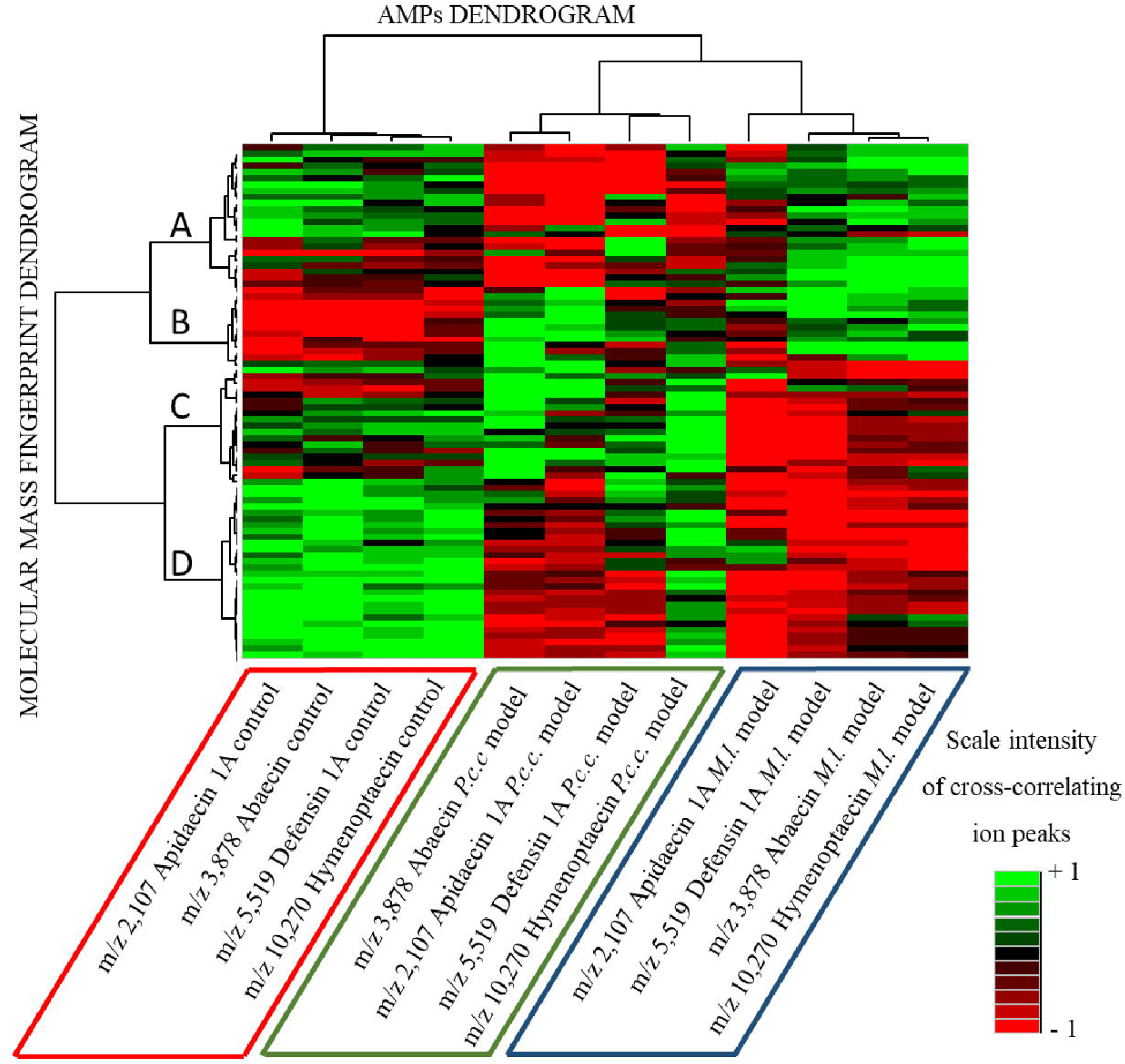
Heat-map of four antimicrobial peptides (AMPs) from *Apis mellifera*, Apidaecin 1A, Abaecin, Defensin 1A and Hymenoptaecin correlating with the MALDI MS fingerprints (102 *m*/*z*) of the hemolymph samples. Per-peak MALDI MS correlation in standard mode between the AMP molecular ions of Apidaecin 1A (*m*/*z* 2,107), Abaecin (*m*/*z* 3,878), Defensin 1A (*m*/*z* 5,519) and Hymenoptaecin (*m*/*z* 10,270), and the MALDI MS fingerprints of the biological model following an experimental infection with either *Micrococcus luteus* (*M. l.*) in blue, or *Pectobacterium carotovorum* subs. *carotovorum* (*P. c. c.*) in green, and the control experiment (non-experimentally infected bees, in red). Each rectangle in the heat map dendrogram represents the abundance level (scale from −1 to +1 from the lowest in red to the highest in green, respectively) of the area of each AMP cross-related with each molecular ion from the fingerprint.

Regarding the control condition, the four AMPs were found to be positively correlated with the molecular ion markers of the hemolymph MFPs of clades A, C and D and negatively correlated with markers of clade B. In the *P. carotovorum* subsp. *carotovorum* model, the same four AMPs were positively correlated with the MFPs of clades B and C and negatively with clade A and D markers, except for Hymenoptaecin, which exhibited positive and negative correlations with clade D. In the *M. luteus* model of infection, each of the AMPs was predominantly positively correlated with the molecular clades A and B, and negatively with clades C and D. These correlations show complementary molecular signatures in the three experimental models. Discrete dynamic molecular patterns are modulated and correlated to the immune status of the bees, allowing us to discriminate infected from non-infected bees and the type of infection

### Machine-learning as the first reported computational model to recognize and classify experimentally infected honey bees based on hemolymph MFPs

Because the proteomic mass spectra of hemolymph samples reflect the immune status of the honey bees, our next goal was to predict honey bee health status based on the bee MALDI-TOF MS MFPs. For that purpose, we decided to build a molecular model based on the MFPs of hemolymph samples, by using a machine-learning algorithm, the Genetic Algorithm (GA). The GA classifier generated a set of discriminating peaks that recognized and classified hemolymph according to the biological model (honey bees challenged with *P. carotovorum carotovorum* or *M. luteus* and non-experimentally infected honey bees). These discriminating peaks form a barcode model and define the strength of this model through its recognition capability. The performance of the classifier barcode model was evaluated through internal cross validation by iterative reclassification of a set of spectra equal to half of the total number of spectra included in the model. For each of the ten iterations performed, a new set of spectra was chosen randomly through an automated internal process.

In an initial approach, we restricted our experimental infection *to M. luteus* as the Gram-positive strain and to *P. carotovorum* subsp. *carotovorum*. While this is, to our knowledge, the first time such a computational model has been applied to the classification of bacterial-infected honey bees, machine-learning algorithms have been used previously in other biological subjects. For example, MALDI MS has been successfully used to build a proteomic mass spectra database of different honeys and their MFPs to identify their geographical origin (51). As another example of application, an experimental model of male chicken fertility was designed to perform on-cell direct proteomic fingerprinting by MALDI MS and demonstrated the capability of the GA classifier to build a predictive model to classify chicken sperm fertility (52).

In the present study, using GA, based on the individual hemolymph spectra of a cohort of 22 controls and 23 honey bees challenged with *M. luteus* or 20 honey bees challenged with *P. carotovorum* subsp. *carotovorum*, we identified a set of nine best *m*/*z* molecular ions based on their capability to discriminate the three biological models from each other (Figure 4). Further tests of recognition capability and cross validation of the GA model were assessed by using the MFPs from the same sample cohort. Considering the standard deviation and the 95% confidence interval of these nine molecular ions, weight indexes were calculated to rank the nine molecular signatures from the most discriminant molecular ion (*m*/*z* 3,348.17, weight of 6.24) to the least discriminant one (*m*/*z* 5,603.01, weight of 1.97). Moreover, we rated the accuracy of the GA classifier model following two distinct data processings. On the one hand, the classifier calculated the recognition capability by matching the MALDI MS spectra described above against their respective biological models. Therefore, we were able to re-assign hemolymph spectra derived from the control and the *M. luteus* biological models (score of recognition 100%) and for the *P. carotovorum* subsp. *carotovorum* model (score of 93.75%). Overall, performance recognition of the classifier reached 97.92%. On the other hand, internal cross validation scores were calculated for each biological model. To perform this cross validation, the same individual hemolymph spectra from each biological model were randomized and reassessed for successful matching in a batch mode of analysis by the classifier using solely the set of the nine molecular ion markers. The cross validations of the classifier were at 91.51%, 94.40% and 89.87% for the control, *M. luteus* and *P. carotovorum* subsp. *carotovorum* biological models, respectively, giving an overall validation of 91.93% (see Fig. 4).

**Figure 4:**
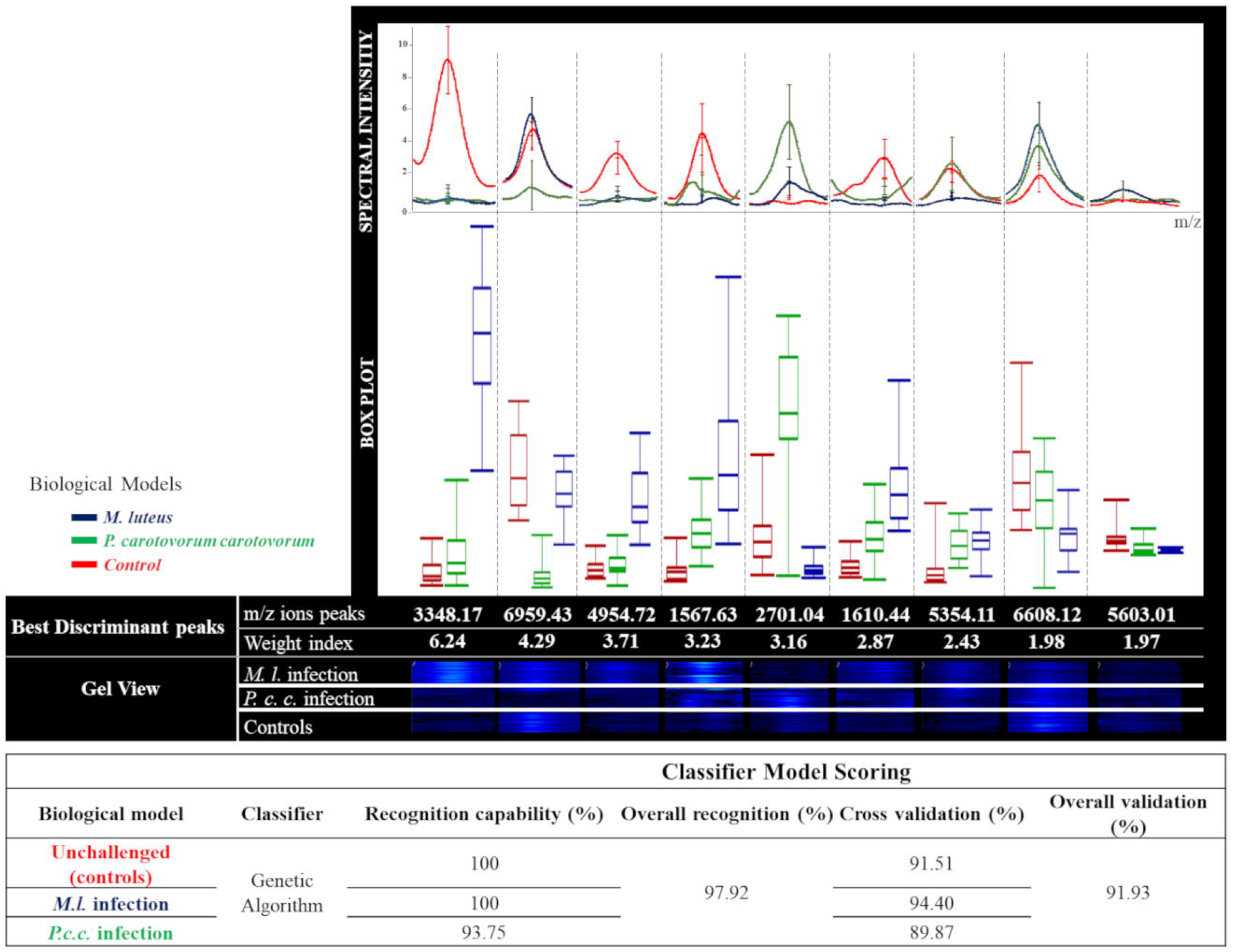
Genetic Algorithm-based classifier used to discriminate non-experimentally infected (Control, red) bees from experimentally infected ones with *Micrococcus luteus* (*M. l.*, in blue) or *Pectobacterium carotovorum* subsp. *carotovorum* (*P. c. c.*, in green). Nine molecular ion peaks determined by the computational model and ranked according to their weight indexes were found as the best discriminative features of the hemolymph samples on the basis of statistical criteria (Standard deviation determined for each curve representing the molecular ions of the model and the box plots showing the first and third interquartile range with line denoting the median and whiskers encompassing 95% of the individuals). The spectral gel view represents the intensity of each of the discriminative peaks represented by their weight index and the *m*/*z* values (Da) found within the individual spectra of the biological models.

A new set made of 26, 10 and 35 MALDI MS spectra of hemolymph from control, *M. luteus* and *P. carotovorum* subsp. *carotovorum* biological models respectively was submitted for the to the GA classifier and classified individually (Table 1 and Table S2). Among the 26 control spectra, 16 were correctly classified, three were classified in *P. carotovorum* subsp. *carotovorum* and three in *M. luteus* models. Four spectra were found as invalid spectra because of the recalibration step. This result was caused by weaker intensities of the molecular fingerprints causing the ion mass recalibration to fail. Regarding the *M. luteus* infectious model, 10 spectra were subjected to the classifier. Seven were correctly classified, two were considered as control and one as belonging to the *P. carotovorum* subsp. *carotovorum* model. No spectrum was deemed invalid. Considering the *P. carotovorum* subsp. *carotovorum* biological model of infection, from the 37 spectra, 13 were correctly classified, one matched to the control, three to the *M. luteus* biological models and 20 to the invalid spectra category. These 20 spectra were qualified as invalid due to noisy mass spectra (intensities of the nine peaks not sufficient to pass the classification) or to a failure in properly calibrating the mass spectra. Given these results, we calculated the performance of the classifier for each of the biological model (see details in Table 2). The GA algorithm achieved 80 % to 90 % accuracy discriminating thus the three biological models. The sensitivity (true positive) and the specificity (true negative) of the GA classifier model were calculated for the three biological models. The model scored at least 70% of sensitivity and at least 84 % of specificity. As detailed in the Table 2, the highest sensitivity was observed for the *P. carotovorum* subsp. *carotovorum* model (76.47 %) and the highest specificity for the control model (95.16 %). Based on the sensitivity and the specificity, we calculated the informedness indexes and the positive-negative stratum-specific likelihood ratio (abbreviated +LR, −LR) which inform about how predictive the classifier model is and its performance as a diagnostic tool respectively. As reported in the Table 2, the three biological models scored indexes within the range [−1;1] with −1 as incorrect model predictions, 1 as maximum of correct predictions). The calculated informedness indexes for the control, the *P. carotovorum* subsp. *carotovorum* scoring 0.68 and 0.64 respectively and for the *M. luteus* (0.55) demonstrated the model was a good predictor. Regarding +LR and −LR, both parameters were calculated. The +LR, which required scores over 1 to be significant were found equal to 15.02; 4.55 and 6.12 for the control, the *M. luteus* and the *P. carotovorum* subsp. *carotovorum* models respectively (Table 2). This result demonstrated a good probability that our GA model classified positively the spectra against the biological models. The −LR, which required scores as close as possible to 0 to be significant were found equal to 0.28; 0.35 and 0.27 for the control, the *M. luteus* and the *P. carotovorum* subsp. *carotovorum* models respectively (Table 2). This result demonstrated the weak probability to missclassify the cohort of hemolymph spectra through the GA classifier. In addition, we determined the false discovery rate (q-value) and the false positive rate (p-value) for each of the three biological models. The lowest q-value was of 0.158 and concerned the control model while *P. carotovorum* subsp. *carotovorum* and *M. luteus* models harbored highest values (0.235 and 0.461 respectively) denoting a better capability of the classifier to classify unkown spectra within the control model followed by the *P. carotovorum* subsp. *carotovorum* and *M. luteus* models respectively. Regarding the p-values, the control model harbored 0.0484 while *P. carotovorum* subsp. *carotovorum* and *M. luteus* models scored 0.125 and 0.154. Based on this statistical parameter, the classifier shares the same conclusion as obtained with the q-values regarding the algorithm’s performance to classify properly the spectra.

To summarize, the computational model significantly discriminated control from infected honey bees and *M. luteus* from *P. carotovorum* subsp. *carotovorum* infection as well on the basis of the hemolymph mass fingerprints.

### MALDI-TOF MS BeeTyping as an effective molecular method to discriminate honey bees infected with different *Serratia* species

We assessed the performance of MALDI BeeTyping to discriminate infection at the species level, in particular between two *Serratia* species: *S. marcescens* isolated from a naturally infected honey bee (*Sm*BIOP160412, Lab. collection) and a reference strain of *S. entomophila* (see dendrogram, Figure 1). We found that the MFPs of the hemolymph samples collected from honey bees infected by *S. entomophila* and *S. marcescens* presented significant molecular differences (Figure 5A).

**Figure 5:**
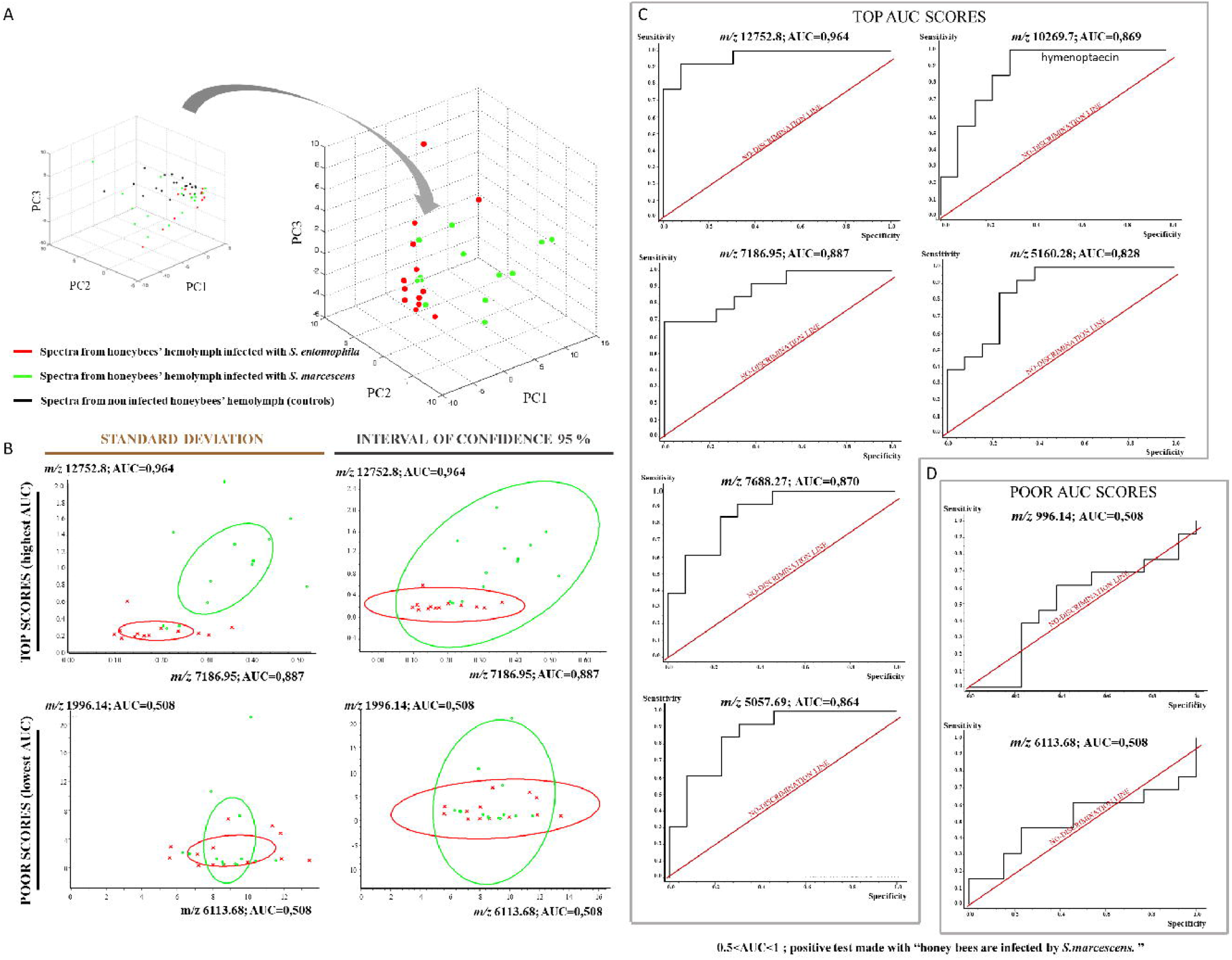
Differential PCA-based analysis of hemolymph fingerprints following infection with two *Serratia* strains, *Serratia marcescens* (*S. m.*) isolated from a naturally infected *Apis mellifera* and *S. entomophila* (*S. e.*) and statistical relevance of predictive markers. The differential fingerprinting and PCA analysis discriminated the non-experimentally infected bees (control in black) and the experimentally infected groups (*S. e.* in red or *S. m*. in green) with n=13 bees per group (**A**). The detected peaks in the differential analysis with the highest and lowest discriminant scores were assessed statistically by measuring the standard deviation and the 95% confidence interval. (**B**). The most interesting peaks classified through the Receiver Operating Characteristics (ROC) curves are shown using the Area Under Curve (AUC) calculation (**C**). The biological model used as the positive class was the experimental *S. m.* infection and the sensibility (True Positives) and specificity (False Positives) parameters were determined for all calculated peaks. Eight markers defined by their *m*/*z* values (Da) were found as the best predictive markers (AUC>0.8) for discriminating honey bee infections (*S. e.* or *S. m.*). Conversely, the two least discriminant peaks had an irrelevant AUC (◻ 0.5) with ROC curves fitting the non-discriminant line of the statistical test (**D**).

Moreover, by using the two best discriminant molecular ions (*m*/*z* 12,752.8 and *m*/*z* 7,186.95), revealed by the PCA analysis, we could differentiate the hemolymph spectra of bees infected either with *S. entomophila* or *S. marcescens Sm*BIOP160412. In contrast, the two weaker discriminant markers *m*/*z* 1,996.14 and *m*/*z* 6,113.68 failed to discriminate the two types of spectra resulting from the two *Serratia* species (Figure 5B).

To further evaluate and rank the measured ion markers within the MFPs based on their capability to discriminate the two *Serratia* species, we performed a receiver operating curve (ROC) analysis to highlight eight *m*/*z* ion markers (12,752.8; 7,186.95; 7,688.27; 10,269.8 (Hymenoptaecin); 5,057.69; 5,160.28) with AUC scores between 0.8 and 1 in sensitivity (Figure 5C). The Apidaecin, Abaecin and Defensin were also checked for their capability to discriminate the two *Serratia* species (Figure S1). Based on the ROC test, Abaecin and Defensin were also capable to discriminate, to some extent, the two *Serratia* species (see Figure S1) (AUC of 0.739 and 0.639, respectively). However, like the two markers *m*/z 1,996.14 and 6,613.68, Apidaecin (AUC=0.520) revealed to be a poor discriminant (Figure 5D). Hence, based on these results, it seems possible to discriminate infection by *S. marcescens* from *S. entomophila* in honey bees using computational modelling. To our knowledge, this is the first report on the feasibility of using MALDI MS as an MFP-based method capable of discriminating hemolymph molecular response to systemic infections induced by two different bacterial species of the same genus. *S. marcescens* is known to be a commensal bacterium present in low abundance in the gut of honey bees. By studying the pathogenicity of different strains of *S. marcescens* through two routes of *in vivo* exposure (oral and direct injection into the hemolymph), Raymann et al (46) found that expression of the four honey bee AMPs Abaecin, Defensin, Hymenoptaecin and Apidaecin did not differ between infected and non-experimentally infected control honey bees. These results support the idea that markers other than AMPs need to be identified and monitored to efficiently discriminate bacterial infections in honey bee hemolymph. As we demonstrated, the correlation of the molecular fingerprints and the AMPs in hemolymph allowed us to discriminate the three different models. In order to determine how specific were the nine markers of the classifier to the control, *M. luteus* and *P. carotovorum* subsp. *carotovorum* biological models, we tested the classification of the hemolymph mass fingerprints of honey bees infected using the two *Serratia* strains. We submitted 13 MALDI mass fingerprints of hemolymph from infected bees with *S. entomophila* to the classifier. One was classified as control, seven as fitting with the *P. carotovorum* subsp. *carotovorum* model, and one fitting with the *M. luteus* model. The classifier excluded four hemolymphs’ fingerprints because of noisy spectra signal or invalid mass recalibration. We also submitted 19 MALDI mass fingerprint of hemolymph from infected bees with *S. marcescens* to the classifier, which identified two as control, five as fitting with the *P. carotovorum* subsp. *carotovorum* model, and four fitting with the *M. luteus* model. The classifier excluded eight hemolymphs’ fingerprints for the same reasons as above. Interestingly, the majority of the classified spectra from both, the *S. entomophila* and *S. marcescens* models matched with the *P. carotovorum* subsp. *carotovorum* model. It is particularly interesting as these three bacteria are Gram-negative. Nevertheless, some spectra matched against the control and the *M. luteus* models. Taking altogether, these results suggest that the BeeTyping approach generates specific molecular barcodes defined accordingly to biological models.

## Conclusion

Along with most relevant, technically feasible and primary observation-based health status indicators highlighted by EFSA’s HEALTHY-B, MALDI-MS BeeTyping, a method derived from the biotyping approach used routinely in clinical microbiology, analyze the downstream responses to stressors through the matured effector molecules circulating in the hemolymph. These effectors include the products of selected immune genes (i.e. genes coding for Apidaecin, Defensin, Hymenoptaecin, and Abaecin) and other molecular mass fingerprints of stress that we are under characterization through a proteomic approach. Our approach of MFPs by MALDI-MS BeeTyping, is a cutting-edge analytic method that may complement and address some limitations issued of the HEALTHY-B toolbox by establishing robust, effective, sensitive and a comprehensive technology for profiling and deciphering, at the individual level, the honeybee health parameters including its immunity stage with regards to bacterial stressors. Moreover, as a robust and sensitive molecular approach, MALDI BeeTyping has several advantages over other molecular biology techniques and visual observations, such as (i) the use of a drop of hemolymph allowing to keep the rest of the body for complementary molecular measurements such as PCR, (ii) a very simple and fast sample preparation, (iii) a short processing time (data acquisition and processing), (iv) low consumable costs, and (v) a user friendly workflow that can be standardized and automated for cost-effective high throughput use. We believe that future developments of MALDI BeeTyping could improve monitoring of honey bee health upon exposure to other biotic or abiotic stressors, the quality control and the origin traceability of apiary products based on molecular markers fingerprinting. Based on specific proteomics signatures, MALDI BeeTyping could bring out a novel analytical tool for early diagnosis of honey bees parasited with *Nosema* species, *Varroa destructor* and infected or not with deformed wing virus or acute bee paralysis virus. We aim at developing the BeeTyping strategy for early diagnosis of honey bees health disorders.

## Material & Methods

The BeeTyping strategy relies on a workflow divided into four major steps summarized in Table S2 and described in this section.

### Biological models

#### Bacterial strains

To generate biological models of infection, we used the Gram-negative strains *Pectobacterium carotovorum* subsp. *carotovorum* 15 (formerly *Erwinia carotovora carotovora* 15 CFBP2141, generous gift from Bruno Lemaitre, EPFL Switzerland), *Serratia entomophila* (Institut Pasteur, CIP102919) and a *Serratia marcescens* strain (*Sm*BIOP160412, our laboratory collection) isolated within the haemocoel from a naturally infected *Apis mellifera* honey bee collected in the field, and the Gram-positive *Micrococcus luteus* (ATCC 4698). Bacteria were cultured in Luria Bertani (LB) medium overnight at 32°C.

#### Bacterial strain identification by MALDI biotyping

The *Pectobacterium carotovorum* subsp. *carotovorum*, *Serratia marcescens*, *S. entomophila* and *Micrococcus luteus* strains were identified following Bruker’s recommendations. Briefly, one isolated colony of bacteria was spread onto a MALDI plate dedicated for microorganism identifications (MALDI Biotarget 48 polished steel) and mixed with 1μ L of Alpha-Cyano-4-hydroxycinnamic acid (4-HCCA) MALDI matrix. Spectra were recorded using the MALDI-TOF MS AutoFlex III instrument and the associated materials, chemicals and software package used for MALDI biotyping were all from Bruker Daltonik (Germany) using the standard method (pre-processing step for which the lower mass was set at 2,000, with a resolution of 5, and a compressing factor of 1). The smoothing frame size was 20Da and the search window was 10Da with three runs for the baseline subtraction. For the peak-picking, the maximum number of peaks was set at 200, with a threshold of 0.0045. The method of peak-picking was based on peak fitting using the Gauss profile. The recorded spectra were matched against the dedicated database MBT Compass 4.1, build 70. The obtained gel spectra for the identified bacteria and their corresponding dendrograms were built under MBT Compass Explorer 4.1 using the standard method of identification. For external calibration of the mass spectrometer, a mix of 1μ L of bacterial test standard proteins (BTS) covering the entire mass range (*m*/*z* 2,000-20,000) of the acquisition method was analyzed using the same protocol.

##### Experimental infection of the honey bees

Experimental infections were performed on newly-emerged honey bee workers (less than 12h old). To design the computational analyses, a training set of spectra was built using non experimentally infected (unpicked control) bees and bees infected with either *Pectobacterium carotovorum* subsp. *carotovorum* 15, *M. luteus, S. entomophila* or the isolated *Serratia marcescens Sm*BIOP160412 strain. Infections were performed by pricking honey bees individually in the anterior lateral thorax (spiracle) using a fine needle (Fine Science Tools, Germany) dipped into a freshly concentrated culture pellet of live bacteria. All honey bees (experimentally infected and controls) were placed for 24h at room temperature in dedicated small cages and fed *ad libitum* with sugar syrup (Invertbee from SARL Isnard, France) containing fructose (36%), dextrose (30%), saccharose (31%), maltose (1.5%) and other sugars (1.5%). Hemolymph was collected from the dorsal side of the abdomen, using pulled glass capillaries (Sutter Instrument Corp, Novato, California). The collected hemolymph was immediately transferred into a chilled LoBind Protein microtube (Eppendorf, Germany) pre-coated with Phenylthiourea and Phenylmethylsulfonyl fluoride (both from Sigma Aldrich, France) to prevent melanization and proteolysis, respectively. The hemolymph samples were stored at −20°C until use.

### Molecular mass fingerprints by MALDI MS

#### Data acquisition

Each individual hemolymph sample was analyzed with the Bruker AutoFlex™ III. The molecular mass fingerprints (MFP) were acquired following the Bruker Biotyper^®^ recommendations (matrix, method of sample deposition and detection) with minor adjustments. Briefly, the hemolymph samples were 10-fold diluted in acidified water (0.1% trifluoroacetic acid, Sigma Aldrich, France) and 0.5μL of a given sample was mixed with 0.5μL of 4-HCCA (Sigma Aldrich, France) on a MALDI MTP 384 polished ground steel plate (Bruker Daltonik). Following co-crystallization of the hemolymph spots with the matrix droplet, MALDI MS spectra were recorded in a linear positive mode and in an automatic data acquisition using FlexControl 4.0 software (Bruker Daltonik). The following instrument settings were used: 1.5kV of electric potential difference, dynamic range of detection of 600 to 18,000 Da, 69% of laser power, a global attenuator offset of 46% with 200Hz laser frequency, and 2,000 accumulated laser shots per hemolymph spectrum with a raster of random walk set to 50. The linear detector gain was setup at 1.82kV with a suppression mass gate up to *m*/*z* 600 to prevent detector saturation by clusters of the 4-HCCA matrix. The pseudo-molecular ions desorbed from the hemolymph were accelerated under 1.5kV. An external calibration of the mass spectrometer was performed using a standard mixture of peptides and proteins (Peptide Standard Calibration II and Protein Standard Calibration I, Bruker Daltonik) covering the dynamic range of analysis.

#### Data post-processing and statistical analyses

The MALDI-MS datasets were imported into the ClinProTools™ 2.2 Software (Bruker Daltonik) for post-processing and statistical analyses. All of the recorded spectra were processed with a baseline subtraction and spectral smoothing followed by an internal recalibration step with exclusion of null and/or “non-recalibratable” spectra. The total averaged spectra were calculated based on a signal over noise ratio equal to 5 for peak-picking and area calculations. The irrelevant spectra that did not pass the required signal intensity and resolution were excluded from any integration into the MALDI-MS computational model designed to match the biological models of honey bee infections. A post-processing step involving spectral normalization of all calculated peak area was performed with ClinProTools™ software prior to statistical analysis (95% confidence interval, standard deviation and Principal Component Analysis-PCA).

#### Hierarchical Clustering, heat maps and ROC curves

The total number of spectra used to design the computational models were normalized and subjected to PCA and unsupervised hierarchical clustering analysis to measure distances between spectra. This analysis was used to determine Euclidean distances (based on PCA results with a reduced dimension limited to 70% and 95% of the total explained variance). The molecular correlation between four antimicrobial peptides (AMPs) known from the honey bee [Apidaecin 1A at *m*/*z* 2,107 (Uniprot entry A0A088AIG0), Hymenoptaecin at *m*/*z* 10,270 (Uniprot entry Q10416), Abaecin at *m*/*z* 3,878 (Uniprot entry P15450) and Defensin 1A at *m*/*z* 5,519 (Uniprot entry P17722)]; and mass fingerprints (MFP) of the three biological models of infection were calculated and represented with a heat map. The receiver operating characteristic (ROC) analyses were built using the ClinProTools™ program and the heat maps, using the OMICs add-on module provided by the XLSTAT program (interquartile threshold value of 0.25).

#### Computational-based algorithm & machine learning model

In the scope of delivering a barcode model capable of discriminating infected from control honey bees, a training set of spectra was established by fingerprinting the corresponding hemolymph samples using MALDI MS. Series of individual spectra were recorded from 22 controls, 23 honey bees challenged with *M. luteus*, 20 with *P. carotovorum* subsp. *carotovorum* 15, and an equal number of 15 spectra from bees challenged with *S. entomophila* and *S. marcescens Sm*BIOP160412. Data clustering (optimal spectral separation combined with the determination of a fixed number of peaks within the training set) was performed using the Genetic Algorithm (GA) with the ClinProTools™ software. The GA parameters were as follow: a maximum of 10 peaks harboring the greatest weight was selected and included in the model. A number of 50 generations (iterative algorithm searching) was chosen to achieve this maximum of peaks. The k-nearest neighbor parameter, which is a key parameter of artificial intelligence used in supervised machine learning, was set at 3.

#### External validation of the barcode model and classification of unknown spectra

In order to assess the capability of the GA classifier to recognize the infected bees from the control group, a new set of hemolymph MS spectra, never processed in the classifier model, was used to perform an external validation. This experimental set of honey bees included the three biological models; 26 controls, 37 infected honey bees with *P. carotovorum* subsp. *carotovorum* 15 and 10 with *M. luteus*. By submitting those hemolymph spectra to the classifier resulted in counting the correctly classified spectra, and also the mismatched and the invalid ones. In order to assess the performance of our classifier model, accuracy, sensitivity, specificity, informedness, specific-positive and negative likelihood ratios, false discovery rate (q-value) and false positive rate (p-value) were calculated. The accuracy, which informs on how efficient the model is, was calculated according to Wang *et al.* (53). Sensitivity scores real positive cases that are correctly predicted positive by the model and the specificity scores the opposite i.e. the real negative cases that are correctly predicted negative. Informedness scores the probability that a prediction (e.g. result of a machine-learning model to classify one condition against the others) is informed regarding to the tested condition *versus* odds. Informedness helps to make diagnosis decision. Sensitivity, specificity and informedness were determined as previously described (54). The specific-positive and -negative likelihood ratios (abbreviated +LR and −LR) classically used in diagnostic testing with multiple classes informs on how likely the results from the classifier model will match the condition. +LR gives the change in the odds of satisfying the condition (fitting to the biological models), given a positive test result and −LR, the change in the odds of satisfying the condition when the test comes negative. +LR ranges from zero to infinity. With +LR values between zero and one, there is a weak probability that the test matches the condition. If the ratio equals to one, then the test lacks diagnostic value and if the ratio is >1, then the test increases the probability to match correctly with the condition. Regarding −LR, the closer to zero the value is, the more informative the test is (55).

## Acknowledgements

We would like to thank Dr. Bruno Lemaitre from the Ecole Polytechnique Fédérale de Lausanne – (EPFL, Switzerland) and the Institut Pasteur de Paris for their generous gifts (*Micrococcus luteus* strain and *Serratia entomophila* respectively). We are also thankful to Paul-Arthur Sol and Erwan Weber for their contribution during their Master 2 training. Financial support was provided by FEAGA (FranceAgriMer), N°14-04R and from the Association Plateforme BioPark of Archamps from its Research & Development budget.

---

Table 1: External validation of the genetic algorithm classifier model using a new set of hemolymph spectra.

Fifty-seven hemolymph samples were collected individually from the biological models *M. l.*, *P. c. c.* and control prior to being fingerprinted. The spectra were submitted to the GA-based computational model in order to assess classifier performance.

Table 2: Assessment of the Genetic Algorithm classifier performance.

Based on the result of the external validation, the performance of the GA-based classifier model was assessed for each of the biological models by calculating the accuracy, the sensitivity, the specificity, the specific-positive and -negative likelihood ratios (all five expressed as percentage), informedness, p-value and q-value.

Table S1: Peak-to-peak correlation scores of the four Antimicrobial peptides (abaecin, apidaecin, defensin and hymenoptaecin) with the mass fingerprint of hemolymph across the three biological models (non-experimentally infected/control, *M. l.* for Micrococcus luteus infection and *P. c. c.* for *Pectobacterium caratovorum subsp. carotovorum 15* infection)

Table S2: Results for the external validation of the genetic algorithm-based classifier

Figure S1: Assessment of ROC curves of Apidaecin, Abaecin and Defensin to discriminate *S. marcescens*- from *S. entomophila*-infected honey bees.

Figure S2: BeeTyping workflow for machine learning data-driven analysis of honey bee infections. The methodological approach relied on four main steps addressing major tasks.

Step 1: Sampling of unchallenged bees (controls) and experimental infection obtained by pricking honey bees with live strains of *P. c. c.* and *M. l‥* Step 2: Individual hemolymph collections followed by MALDI-TOF MS molecular mass fingerprinting, and strain identification by MALDI biotyping. Step 3: Multi-stage processing of MALDI MS fingerprints including recalibration, peak picking, normalization and statistical calculation of individual MS spectra through Principal Component Analysis (PCA) for revealing differential molecular patterns across infection groups. Step 4: Genetic algorithm-based computational model for recognition and classification of honey bee infection using PCA discriminant analysis. Barcodes were built following the molecular fingerprints that discriminate control bees from bees infected either with *M. l.* or with *P. c. c.*.

## References

1. Biesmeijer JC, et al. (2006) Parallel declines in pollinators and insect-pollinated plants in Britain and the Netherlands. Science 313(5785):351–354.

2. Potts SG, et al. (2010) Global pollinator declines: trends, impacts and drivers. Trends in ecology & evolution 25(6):345–353.

3. Aj V (2013) Threats to an ecosystem service: pressures on pollinators. Front Ecol Environ 11(5):251–259.

4. Koh I, et al. (2016) Modeling the status, trends, and impacts of wild bee abundance in the United States. Proceedings of the National Academy of Sciences of the United States of America 113(1):140–145.

5. Goulson D, Nicholls E, Botias C, & Rotheray EL (2015) Bee declines driven by combined stress from parasites, pesticides, and lack of flowers. Science 347(6229):1255957.

6. Breeze TD, et al. (2014) Agricultural policies exacerbate honeybee pollination service supply-demand mismatches across Europe. PloS one 9(1):e82996.

7. Smith KM, et al. (2013) Pathogens, pests, and economics: drivers of honey bee colony declines and losses. EcoHealth 10(4):434–445.

8. Dainat B, Vanengelsdorp D, & Neumann P (2012) Colony collapse disorder in Europe. Environmental microbiology reports 4(1):123–125.

9. Neumann P. CN (2010) Honey bee colony losses. Journal of apicultural research 49:1–6.

10. Steinhauer N, et al. (2018) Drivers of colony losses. Current opinion in insect science 26:142–148.

11. Mullin CA, et al. (2010) High levels of miticides and agrochemicals in North American apiaries: implications for honey bee health. PloS one 5(3):e9754.

12. Wu JY, Smart MD, Anelli CM, & Sheppard WS (2012) Honey bees (Apis mellifera) reared in brood combs containing high levels of pesticide residues exhibit increased susceptibility to Nosema (Microsporidia) infection. Journal of invertebrate pathology 109(3):326–329.

13. Clermont A, Eickermann M, Kraus F, Hoffmann L, & Beyer M (2015) Correlations between land covers and honey bee colony losses in a country with industrialized and rural regions. The Science of the total environment 532:1–13.

14. Smart M, Pettis J, Rice N, Browning Z, & Spivak M (2016) Linking Measures of Colony and Individual Honey Bee Health to Survival among Apiaries Exposed to Varying Agricultural Land Use. PloS one 11(3):e0152685.

15. Beyer M, et al. (2018) Winter honey bee colony losses, Varroa destructor control strategies, and the role of weather conditions: Results from a survey among beekeepers. Research in veterinary science 118:52–60.

16. Traver BE, Feazel-Orr HK, Catalfamo KM, Brewster CC, & Fell RD (2018) Seasonal Effects and the Impact of In-Hive Pesticide Treatments on Parasite, Pathogens, and Health of Honey Bees. Journal of economic entomology 111(2):517–527.

17. Desai SD & Currie RW (2016) Effects of Wintering Environment and Parasite-Pathogen Interactions on Honey Bee Colony Loss in North Temperate Regions. PloS one 11(7):e0159615.

18. Di Pasquale G, et al. (2013) Influence of pollen nutrition on honey bee health: do pollen quality and diversity matter? PloS one 8(8):e72016.

19. Alaux C, Dantec C, Parrinello H, & Le Conte Y (2011) Nutrigenomics in honey bees: digital gene expression analysis of pollen’s nutritive effects on healthy and varroa-parasitized bees. BMC genomics 12:496.

20. Dolezal AG & Toth AL (2018) Feedbacks between nutrition and disease in honey bee health. Current opinion in insect science 26:114–119.

21. Sinpoo C, Paxton RJ, Disayathanoowat T, Krongdang S, & Chantawannakul P (2018) Impact of Nosema ceranae and Nosema apis on individual worker bees of the two host species (Apis cerana and Apis mellifera) and regulation of host immune response. Journal of insect physiology 105:1–8.

22. Nazzi F, Annoscia D, Caprio E, Di Prisco G, & Pennacchio F (2014) Honeybee immunity and colony losses. Entomologia 2(203):80–86.

23. Flenniken ML & Andino R (2013) Non-specific dsRNA-mediated antiviral response in the honey bee. Plos One 8(10):e77263.

24. Zhang Y, Liu X, Zhang W, & Han R (2010) Differential gene expression of the honey bees Apis mellifera and A. cerana induced by Varroa destructor infection. Journal of insect physiology 56(9):1207–1218.

25. Di Prisco G, et al. (2013) Neonicotinoid clothianidin adversely affects insect immunity and promotes replication of a viral pathogen in honey bees. Proceedings of the National Academy of Sciences of the United States of America 110(46):18466–18471.

26. (2016) E (2016) Assessing the health status of managed honeybee colonies (HEALTHY-B): a tool box to facilitate harmonised data collection. EFSA Journal 14(10):1–248.

27. Meikle W, Holst N (2015) Application of continuous monitoring of honeybee colonies. Apidologie 46:10–22.

28. Gil-Lebrero S, et al. (2016) Honey Bee Colonies Remote Monitoring System. Sensors 17(1).

29. Becher MA, et al. (2018) Bumble-BEEHAVE: A systems model for exploring multifactorial causes of bumblebee decline at individual, colony, population and community level. The Journal of applied ecology 55(6):2790–2801.

30. Karas M (2012) A history of european mass spectrometry (IM Publications LLP).

31. Spinali S, et al. (2015) Microbial typing by matrix-assisted laser desorption ionization-time of flight mass spectrometry: do we need guidance for data interpretation? Journal of clinical microbiology 53(3):760–765.

32. De Respinis S, et al. (2014) Matrix-assisted laser desorption ionization-time of flight (MALDI-TOF) mass spectrometry using the Vitek MS system for rapid and accurate identification of dermatophytes on solid cultures. Journal of clinical microbiology 52(12):4286–4292.

33. van Belkum A, et al. (2013) Rapid clinical bacteriology and its future impact. Annals of laboratory medicine 33(1):14–27.

34. Senan S, et al. (2015) Geriatric Respondents and Non-Respondents to Probiotic Intervention Can be Differentiated by Inherent Gut Microbiome Composition. Frontiers in microbiology 6:944.

35. Bourassa L, Susan M, & Butler-Wu (2015) Current and Emerging Technologies for the Diagnosis of Microbial Infections. Method in Microbiology:37–87.

36. Clark AE, Kaleta EJ, Arora A, & Wolk DM (2013) Matrix-assisted laser desorption ionization-time of flight mass spectrometry: a fundamental shift in the routine practice of clinical microbiology. Clinical microbiology reviews 26(3):547–603.

37. Sabatier L, et al. (2003) Pherokine-2 and -3. European journal of biochemistry 270(16):3398–3407.

38. Boulanger N, et al. (2001) Immune response of Drosophila melanogaster to infection with the flagellate parasite Crithidia spp. Insect biochemistry and molecular biology 31(2):129–137.

39. Uttenweiler-Joseph S, et al. (1998) Differential display of peptides induced during the immune response of Drosophila: a matrix-assisted laser desorption ionization time-of-flight mass spectrometry study. Proceedings of the National Academy of Sciences of the United States of America 95(19):11342–11347.

40. Casteels-Josson K, Zhang W, Capaci T, Casteels P, & Tempst P (1994) Acute transcriptional response of the honeybee peptide-antibiotics gene repertoire and required post-translational conversion of the precursor structures. The Journal of biological chemistry 269(46):28569–28575.

41. Casteels P, Ampe C, Jacobs F, Vaeck M, & Tempst P (1989) Apidaecins: antibacterial peptides from honeybees. The EMBO journal 8(8):2387–2391.

42. Ishii K, Hamamoto H, & Sekimizu K (2014) Establishment of a bacterial infection model using the European honeybee, Apis mellifera L. PloS one 9(2):e89917.

43. Hetru C & Bulet P (1997) Strategies for the isolation and characterization of antimicrobial peptides of invertebrates. Methods in molecular biology 78:35–49.

44. Pomastowski P, et al. (2019) Analysis of bacteria associated with honeys of different geographical and botanical origin using two different identification approaches: MALDI-TOF MS and 16S rDNA PCR technique. PloS one 14(5):e0217078.

45. Acosta Muniz C, Jaillard D, Lemaitre B, & Boccard F (2007) Erwinia carotovora Evf antagonizes the elimination of bacteria in the gut of Drosophila larvae. Cellular microbiology 9(1):106–119.

46. Raymann K, Coon KL, Shaffer Z, Salisbury S, & Moran NA (2018) Pathogenicity of Serratia marcescens Strains in Honey Bees. mBio 9(5).

47. Motta EVS, Raymann K, & Moran NA (2018) Glyphosate perturbs the gut microbiota of honey bees. Proceedings of the National Academy of Sciences of the United States of America 115(41):10305–10310.

48. Jiří Danihlík KAMP (2015) Antimicrobial peptides: a key component of honey bee innate immunity Journal of apicultural research 54(2):123–136.

49. Casteels P, Ampe C, Jacobs F, & Tempst P (1993) Functional and chemical characterization of Hymenoptaecin, an antibacterial polypeptide that is infection-inducible in the honeybee (Apis mellifera). The Journal of biological chemistry 268(10):7044–7054.

50. Casteels P, et al. (1990) Isolation and characterization of abaecin, a major antibacterial response peptide in the honeybee (Apis mellifera). European journal of biochemistry 187(2):381–386.

51. Wang J, et al. (2009) Rapid determination of the geographical origin of honey based on protein fingerprinting and barcoding using MALDI TOF MS. Journal of agricultural and food chemistry 57(21):10081–10088.

52. Soler L, et al. (2016) Intact Cell MALDI-TOF MS on Sperm: A Molecular Test For Male Fertility Diagnosis. Molecular & cellular proteomics : MCP 15(6):1998–2010.

53. Wang H.Y., et al. (2018) Application of a MALDI-TOF analysis platform (ClinProTools) for rapid and preliminary report of MRSA sequence types in Taiwan. PeerJ. 7(6):e5784

54. Powers, D.M.W. (2011) Evaluation: From precision, recall and F-measure to ROC, informedness, markedness & correlation. Journal of Machine Learning Technologies. 2(1):37–63

55. McGee, S. (2011) Simplifying Likelihood Ratios. J Gen Intern Med. 17(8):647–650

