## Supplementary material for "A MALDI-MS biotyping-like method to address honey bee health status through computational modelling": Table S1

| | Correlation of Abaecin | | | Correlation of Apidaecin | | | Correlation of Defensin 1 | | | Correlation of Hymenoptaecin | | |
|---|---|---|---|---|---|---|---|---|---|---|---|---|
| **Mass Fingerprint peaks m/z** | **Control model** | ***P. c. c.* model** | ***M. l.* model** | **Control model** | ***P. c. c.* model** | ***M. l.* model** | **Control model** | ***P. c. c.* model** | ***M. l.* model** | **Control model** | ***P. c. c.* model** | ***M. l.* model** |
| **1514.95** | -0.263 | -0.5369 | -0.1494 | -0.3104 | -0.6593 | -0.2073 | -0.2232 | -0.4211 | -0.2808 | -0.098 | -0.1234 | -0.2839 |
| **1542.78** | -0.4447 | -0.1948 | -0.0969 | -0.2488 | -0.503 | 0.0215 | -0.3802 | -0.0194 | -0.1337 | -0.2221 | 0.1178 | -0.1848 |
| **1567.22** | -0.0973 | -0.4294 | 0.092 | 0.0117 | -0.5702 | 0.1046 | -0.0044 | -0.4972 | 0.0107 | 0.3196 | -0.185 | 0.0357 |
| **1586.22** | -0.2154 | -0.6346 | -0.3456 | -0.2234 | -0.5372 | -0.1345 | -0.2245 | -0.5312 | -0.3949 | -0.0027 | -0.3275 | -0.4249 |
| **1596.26** | -0.1868 | -0.3929 | -0.2404 | -0.2659 | -0.5325 | -0.1418 | -0.2792 | 0.3882 | -0.255 | -0.3742 | -0.2564 | -0.3065 |
| **1610.26** | 0.2264 | -0.336 | 0.3894 | 0.3253 | -0.1327 | 0.0556 | 0.1845 | -0.0147 | 0.3592 | 0.0844 | -0.1417 | 0.3111 |
| **1614.94** | 0.2396 | -0.7412 | 0.3516 | 0.298 | -0.5554 | -0.261 | 0.169 | -0.0156 | 0.2773 | 0.1706 | -0.3612 | 0.3101 |
| **1667.39** | 0.1946 | -0.0537 | -0.1605 | 0.3411 | -0.0022 | -0.1653 | 0.0782 | -0.0107 | -0.2854 | -0.032 | 0.1684 | 0.0237 |
| **1712.57** | -0.1829 | -0.4057 | 0.1357 | 0.142 | -0.3644 | -0.3259 | -0.1987 | -0.5901 | -0.1122 | -0.2256 | -0.0865 | 0.2035 |
| **1726.54** | -0.1508 | -0.5313 | 0.5808 | 0.0728 | -0.3111 | 0.1413 | -0.2392 | -0.1677 | 0.3544 | -0.1935 | -0.3777 | 0.7548 |
| **1819.03** | 0.3597 | -0.2889 | 0.1684 | 0.462 | -0.1881 | 0.1906 | 0.2841 | -0.2793 | 0.2269 | 0.0894 | -0.2664 | 0.2085 |
| **1837.30** | 0.6735 | 0.5956 | 0.3171 | 0.8859 | 0.5 | 0.5592 | 0.7865 | 0.078 | 0.5133 | 0.4672 | 0.2184 | 0.2508 |
| **1945.20** | 0.2618 | -0.6599 | -0.6558 | -0.0489 | -0.5294 | -0.5259 | 0.1395 | -0.271 | -0.6289 | 0.2585 | -0.4611 | -0.7083 |
| **1954.96** | 0.0898 | -0.8132 | -0.6575 | -0.0459 | -0.6785 | -0.1142 | -0.0367 | -0.2986 | -0.5128 | 0.1602 | -0.4195 | -0.7483 |
| **1993.6** | 0.0162 | 0.3037 | -0.7648 | -0.1341 | 0.2987 | -0.4302 | -0.009 | -0.1751 | -0.5453 | -0.1534 | 0.1442 | -0.8405 |
| **2065.84** | 0.7517 | -0.3536 | 0.6126 | 0.7819 | -0.1073 | 0.4972 | 0.7009 | -0.4565 | 0.6779 | 0.4298 | 0.1931 | 0.6834 |
| **2092.15** | 0.8646 | 0.4053 | 0.6163 | 0.9544 | 0.801 | 0.6061 | 0.8204 | -0.06 | 0.7012 | 0.597 | 0.2724 | 0.6621 |
| **2107.73** | 0.751 | 0.639 | 0.2368 | n.a. | n.a. | n.a. | 0.8004 | 0.2986 | 0.4456 | 0.5578 | 0.3608 | 0.218 |
| **2128.69** | 0.5922 | 0.4382 | 0.487 | 0.8428 | 0.8963 | 0.5136 | 0.5688 | 0.2497 | 0.3387 | 0.5843 | 0.3248 | 0.3746 |
| **2145.49** | 0.6951 | 0.7121 | -0.1057 | 0.9535 | 0.8604 | 0.6514 | 0.781 | 0.2158 | 0.1553 | 0.5403 | 0.3332 | -0.2551 |
| **2248.94** | -0.0604 | -0.5651 | 0.2021 | 0.1529 | -0.4651 | -0.3869 | 0.031 | 0.1061 | -0.1125 | 0.0239 | -0.3386 | 0.0419 |
| **2412.01** | 0.0997 | -0.1497 | -0.0389 | 0.2507 | -0.4576 | -0.694 | 0.0138 | 0.2915 | -0.2137 | -0.0538 | -0.1969 | -0.0575 |
| **2559.79** | -0.1597 | 0.0632 | -0.1582 | -0.2211 | -0.1848 | -0.1145 | -0.3061 | 0.5091 | -0.3626 | -0.1071 | -0.2208 | 0.0333 |
| **2701.32** | -0.3491 | -0.8514 | -0.427 | -0.3099 | -0.7879 | -0.5607 | -0.467 | -0.3422 | -0.5468 | -0.4676 | -0.4669 | -0.351 |
| **2760.25** | 0.8208 | 0.1986 | -0.3021 | 0.8403 | 0.1312 | -0.8234 | 0.8376 | 0.2256 | -0.5328 | 0.6776 | 0.0752 | -0.1336 |
| **2856.29** | -0.07 | -0.0309 | -0.554 | -0.113 | 0.3629 | -0.1898 | -0.0973 | -0.1077 | -0.2702 | -0.1489 | 0.0917 | -0.6361 |
| **2962.41** | -0.1307 | 0.2599 | -0.4371 | -0.3589 | 0.166 | -0.8778 | -0.3559 | -0.1867 | -0.6339 | -0.5307 | 0.2247 | -0.4369 |
| **2983.07** | -0.2385 | 0.1357 | 0.2615 | -0.3454 | 0.4409 | -0.134 | -0.2751 | -0.4102 | 0.3396 | -0.4741 | -0.0411 | 0.2203 |
| **3195.23** | 0.1851 | -0.3578 | -0.5697 | -0.1307 | -0.6777 | -0.815 | -0.0793 | -0.0499 | -0.6515 | -0.1288 | -0.2865 | -0.5438 |
| **3217.82** | 0.1637 | -0.232 | -0.7092 | 0.0424 | -0.6648 | -0.6396 | -0.1484 | -0.0794 | -0.6993 | -0.1473 | -0.2801 | -0.7665 |
| **3235.25** | -0.0044 | -0.6518 | -0.6679 | -0.1576 | -0.6568 | -0.7751 | -0.2316 | -0.625 | -0.6914 | -0.1497 | -0.2818 | -0.6985 |
| **3308.51** | 0.2011 | 0.5002 | 0.3535 | -0.0498 | 0.3493 | 0.1719 | 0.1123 | 0.466 | 0.3401 | 0.0858 | 0.1865 | 0.5634 |
| **3331.80** | 0.1645 | 0.3336 | 0.3159 | -0.0478 | -0.0673 | 0.4771 | 0.1312 | 0.5337 | 0.1394 | 0.1616 | 0.0968 | 0.4425 |
| **3348.18** | 0.0686 | 0.6191 | -0.377 | -0.1104 | 0.2675 | 0.4608 | 0.0407 | 0.3285 | -0.1794 | 0.2352 | 0.1673 | -0.2695 |
| **3478.43** | 0.0054 | 0.1588 | -0.0597 | -0.1231 | -0.1202 | -0.7816 | -0.0485 | -0.0767 | -0.1901 | -0.0339 | 0.5758 | 0.006 |
| **3782.22** | -0.3132 | -0.3768 | -0.2553 | -0.057 | -0.3476 | 0.0668 | -0.2723 | -0.243 | -0.1776 | -0.2557 | 0.1781 | -0.1571 |
| **3878.78** | n.a. | n.a. | n.a. | 0.751 | 0.639 | 0.2368 | 0.8589 | 0.4763 | 0.7761 | 0.6578 | 0.5873 | 0.842 |
| **4014.66** | -0.285 | 0.0119 | -0.3698 | -0.1973 | -0.3035 | -0.1268 | -0.1852 | -0.044 | -0.2272 | -0.2842 | 0.3811 | -0.2526 |
| **4029.84** | -0.2093 | 0.0096 | -0.5287 | -0.2013 | -0.4843 | -0.2582 | -0.1809 | -0.1113 | -0.3437 | -0.149 | 0.0248 | -0.4363 |
| **4135.26** | 0.2936 | 0.5417 | 0.4434 | 0.0418 | 0.7682 | 0.3268 | -0.1266 | 0.5821 | 0.5136 | -0.0346 | 0.2739 | 0.3769 |
| **4190.98** | -0.0479 | 0.127 | -0.3091 | -0.2409 | -0.1302 | -0.1651 | -0.0515 | 0.7047 | -0.308 | -0.0029 | -0.1392 | -0.5359 |
| **4322.85** | -0.246 | 0.4352 | -0.1405 | 0.0425 | 0.6277 | -0.1857 | -0.133 | -0.0612 | -0.1307 | 0.0574 | 0.387 | -0.2298 |
| **4338.89** | -0.2953 | 0.0885 | -0.3637 | -0.0065 | -0.2962 | -0.857 | -0.1665 | 0.2458 | -0.5367 | 0.0274 | 0.297 | -0.4077 |
| **4418.00** | 0.0648 | -0.4791 | 0.353 | 0.091 | -0.6419 | -0.1269 | -0.24 | 0.0279 | 0.1797 | -0.1084 | -0.3342 | 0.3367 |
| **4432.24** | 0.1572 | -0.6524 | 0.194 | 0.2745 | -0.6115 | 0.2857 | -0.1398 | -0.7168 | 0.1391 | 0.0048 | -0.2633 | 0.4296 |
| **4550.31** | 0.0845 | -0.6097 | -0.4411 | -0.0028 | -0.6916 | -0.7409 | 0.1155 | -0.7017 | -0.5819 | 0.1153 | -0.1052 | -0.4069 |
| **4560.25** | 0.2449 | -0.7242 | -0.5995 | 0.2841 | -0.6052 | -0.7219 | 0.1966 | -0.5966 | -0.6522 | 0.3258 | -0.283 | -0.4872 |
| **4673.45** | -0.2256 | -0.6609 | -0.1304 | 0.0358 | -0.6118 | -0.2432 | -0.392 | -0.7283 | -0.2193 | -0.345 | -0.3692 | -0.2948 |
| **4688.41** | -0.2367 | -0.6321 | -0.3968 | -0.2237 | -0.6758 | -0.6606 | -0.3994 | -0.6051 | -0.5364 | -0.3783 | -0.3364 | -0.4756 |
| **4729.28** | 0.0853 | 0.3379 | -0.0401 | -0.115 | -0.0855 | -0.2288 | 0.0489 | 0.6919 | -0.1196 | 0.2434 | 0.0287 | 0.1591 |
| **4954.77** | 0.283 | -0.0528 | 0.0855 | 0.2775 | -0.4003 | 0.2283 | 0.3268 | 0.3769 | 0.1226 | 0.3673 | -0.3377 | 0.3862 |
| **5268.19** | 0.0573 | -0.743 | -0.2612 | 0.2483 | -0.6855 | -0.7974 | 0.0875 | -0.3887 | -0.5636 | 0.0727 | -0.2967 | -0.1636 |
| **5354.47** | 0.076 | -0.0512 | -0.1496 | 0.2032 | -0.2496 | -0.1126 | -0.2081 | 0.1813 | -0.1631 | -0.108 | 0.0236 | 0.113 |
| **5395.51** | 0.0631 | -0.2911 | -0.6199 | -0.1611 | -0.4432 | -0.6304 | -0.1261 | -0.1177 | -0.7238 | 0.0295 | -0.0086 | -0.5978 |
| **5466.30** | 0.8416 | 0.6684 | 0.0832 | 0.6067 | 0.2475 | -0.2753 | 0.8142 | 0.6053 | -0.1306 | 0.6316 | 0.4711 | 0.1359 |
| **5519.26** | 0.8589 | 0.4763 | 0.7761 | 0.8004 | 0.2986 | 0.4456 | n.a. | n.a. | n.a. | 0.7702 | 0.2124 | 0.6595 |
| **5602.67** | 0.1012 | 0.6683 | -0.2371 | 0.212 | 0.2733 | -0.3834 | -0.1986 | 0.4466 | -0.1797 | -0.0824 | 0.6513 | -0.3835 |
| **5658.60** | -0.2062 | 0.3191 | 0.167 | -0.1696 | 0.6069 | -0.2674 | -0.2287 | 0.0832 | 0.2097 | -0.0574 | 0.1872 | -0.0164 |
| **5711.57** | -0.4684 | 0.4493 | 0.7774 | -0.2753 | 0.6602 | 0.2282 | -0.2968 | 0.3525 | 0.7216 | -0.1958 | 0.1153 | 0.5639 |
| **5750.06** | -0.4816 | 0.4009 | 0.3074 | -0.265 | 0.5294 | -0.1215 | -0.4075 | 0.2887 | 0.2125 | -0.1312 | 0.0151 | 0.1236 |
| **5920.96** | -0.379 | 0.1904 | 0.3471 | -0.4251 | 0.3692 | 0.4867 | -0.3143 | 0.0191 | 0.458 | -0.2439 | -0.1779 | 0.227 |
| **5963.55** | -0.5531 | 0.4897 | 0.4334 | -0.5468 | 0.7795 | 0.5388 | -0.4844 | 0.0229 | 0.643 | -0.5199 | 0.03 | 0.3571 |
| **5977.17** | -0.1504 | -0.0733 | 0.3316 | -0.3595 | 0.2972 | 0.198 | -0.1798 | -0.3221 | 0.3089 | -0.3917 | -0.2484 | 0.2374 |
| **6117.70** | -0.004 | 0.5893 | -0.3778 | 0.1489 | 0.7987 | -0.8719 | 0.1306 | 0.5206 | -0.569 | 0.1766 | 0.4836 | -0.3914 |
| **6196.48** | -0.0267 | -0.172 | -0.3549 | 0.0757 | -0.111 | -0.7439 | 0.0728 | 0.0059 | -0.45 | 0.0783 | 0.2285 | -0.3331 |
| **6211.90** | 0.0647 | -0.533 | -0.3475 | 0.2283 | -0.4512 | -0.7986 | 0.1963 | -0.5712 | -0.4756 | 0.1663 | 0.1128 | -0.3052 |
| **6239.86** | -0.006 | -0.3522 | -0.6162 | 0.115 | -0.3635 | 0.0561 | 0.0834 | -0.0992 | -0.473 | 0.0387 | 0.015 | -0.5788 |
| **6377.34** | 0.135 | -0.2026 | 0.1513 | -0.1261 | -0.2669 | -0.0714 | -0.0962 | 0.5127 | 0.1759 | -0.0683 | -0.1101 | 0.2624 |
| **6607.89** | -0.1074 | -0.5358 | 0.117 | -0.3802 | -0.5472 | -0.0971 | -0.1503 | -0.2986 | 0.1428 | -0.1288 | -0.0592 | 0.2541 |
| **6756.47** | -0.1224 | 0.24 | 0.2682 | -0.3549 | -0.0171 | -0.3972 | -0.1387 | -0.2597 | 0.2424 | -0.1552 | 0.1222 | 0.3521 |
| **6923.56** | -0.053 | 0.1236 | -0.3397 | -0.2805 | -0.1322 | -0.8729 | -0.093 | -0.4869 | -0.5261 | -0.1504 | 0.324 | -0.3173 |
| **6958.86** | -0.0489 | 0.7736 | 0.3839 | -0.1573 | 0.8712 | 0.2849 | -0.0272 | 0.4017 | 0.5133 | 0.264 | 0.4716 | 0.6005 |
| **7137.23** | 0.2869 | -0.5244 | -0.4866 | 0.2727 | -0.5736 | -0.5584 | 0.0859 | -0.542 | -0.6482 | 0.2056 | 0.032 | -0.6117 |
| **7184.52** | 0.0457 | -0.427 | -0.2266 | -0.0712 | -0.5298 | -0.0745 | -0.2343 | -0.2924 | -0.277 | 0.0433 | 0.0476 | -0.2382 |
| **7324.52** | -0.1366 | -0.7277 | -0.3517 | -0.1354 | -0.614 | -0.6967 | -0.3035 | -0.7625 | -0.6211 | -0.2298 | -0.2843 | -0.2687 |
| **7339.42** | 0.0613 | -0.7105 | -0.4204 | 0.0636 | -0.5871 | -0.8332 | -0.1104 | -0.7005 | -0.6961 | 0.0132 | -0.1793 | -0.4316 |
| **7382.09** | 0.2143 | -0.5768 | -0.4005 | 0.2371 | -0.5423 | -0.8197 | 0.0792 | -0.7376 | -0.5807 | 0.1272 | 0.0305 | -0.3838 |
| **7689.68** | -0.1777 | 0.3285 | 0.4061 | -0.1875 | 0.0101 | -0.0217 | -0.1572 | 0.6089 | 0.3914 | -0.0428 | 0.0163 | 0.3968 |
| **8190.05** | 0.0684 | 0.4062 | -0.2523 | 0.2523 | 0.0884 | -0.5638 | 0.1007 | 0.172 | -0.5429 | 0.0898 | 0.6296 | -0.3329 |
| **8369.16** | 0.2491 | -0.3498 | -0.4896 | 0.2189 | -0.5334 | -0.5845 | 0.0306 | 0.1791 | -0.5822 | -0.1034 | 0.0107 | -0.4386 |
| **8410.79** | 0.197 | -0.3015 | -0.6571 | 0.016 | -0.5073 | -0.4327 | -0.0956 | -0.3294 | -0.6622 | -0.0087 | 0.0431 | -0.575 |
| **8538.57** | 0.1257 | -0.5299 | 0.3762 | -0.0894 | -0.5994 | 0.268 | -0.0233 | -0.5699 | 0.3338 | 0.1214 | 0.0363 | 0.452 |
| **8762.49** | 0.2172 | -0.304 | -0.4328 | 0.2394 | -0.406 | -0.6101 | 0.288 | -0.5357 | -0.6021 | 0.2717 | 0.1946 | -0.4421 |
| **8830.46** | 0.215 | -0.426 | -0.4821 | 0.054 | -0.5762 | -0.7505 | 0.1384 | -0.1509 | -0.7215 | 0.3227 | 0.0257 | -0.4964 |
| **8948.98** | 0.058 | -0.5111 | -0.6527 | -0.1366 | -0.4438 | -0.7132 | -0.2562 | -0.7656 | -0.7075 | -0.191 | 0.0005 | -0.5714 |
| **9079.42** | -0.0113 | -0.5433 | -0.321 | -0.1037 | -0.5039 | -0.8488 | 0.0194 | -0.5352 | -0.5536 | 0.0082 | -0.0114 | -0.3185 |
| **9267.52** | 0.2699 | -0.4732 | -0.4276 | 0.2333 | -0.6008 | -0.7502 | 0.1152 | -0.4744 | -0.62 | 0.3238 | -0.1002 | -0.3517 |
| **9313.63** | -0.1457 | 0.1094 | 0.048 | -0.0178 | -0.2165 | -0.3137 | -0.1434 | 0.5175 | -0.1114 | -0.0409 | -0.1588 | 0.1662 |
| **9518.22** | -0.123 | -0.4554 | -0.4607 | -0.0518 | -0.4552 | -0.2363 | -0.274 | -0.2982 | -0.4582 | -0.1637 | 0.0565 | -0.4998 |
| **9591.87** | -0.0491 | -0.3688 | -0.348 | -0.134 | -0.3578 | -0.7683 | -0.1607 | -0.3134 | -0.5297 | -0.1525 | 0.0058 | -0.2756 |
| **9683.33** | 0.0489 | -0.3563 | -0.6268 | -0.0535 | -0.4352 | -0.3998 | -0.0198 | -0.0624 | -0.7446 | 0.2162 | 0.1311 | -0.5915 |
| **9966.41** | 0.2029 | -0.3668 | -0.6737 | -0.1881 | -0.5883 | -0.1078 | 0.001 | -0.5473 | -0.5274 | -0.009 | -0.0825 | -0.6319 |
| **10160.88** | 0.4069 | -0.1591 | -0.6773 | 0.0507 | -0.4951 | -0.3489 | 0.2462 | -0.2626 | -0.57 | 0.4238 | 0.3503 | -0.6365 |
| **10270.67** | 0.6578 | 0.5873 | 0.842 | 0.5578 | 0.3608 | 0.218 | 0.7702 | 0.2124 | 0.6595 | n.a. | n.a. | n.a. |
| **10321.78** | 0.2193 | 0.6997 | 0.2225 | -0.089 | 0.2834 | 0.3151 | 0.2987 | 0.3368 | 0.0964 | 0.5285 | 0.8209 | 0.4733 |
| **10393.67** | -0.0722 | 0.8694 | 0.7059 | 0.0641 | 0.7516 | 0.1942 | 0.0684 | 0.5612 | 0.5307 | 0.2782 | 0.6007 | 0.8646 |
| **10408.71** | 0.116 | 0.8456 | 0.6152 | 0.2541 | 0.5596 | 0.0681 | 0.2871 | 0.3972 | 0.3492 | 0.4688 | 0.6673 | 0.8441 |
| **10447.60** | 0.684 | 0.7627 | 0.4654 | 0.5073 | 0.2357 | -0.4744 | 0.7659 | 0.3909 | 0.1028 | 0.8611 | 0.7303 | 0.5931 |
| **10522.45** | 0.3477 | 0.5982 | -0.2858 | 0.1878 | 0.0461 | -0.835 | 0.4597 | 0.0541 | -0.5453 | 0.5277 | 0.6493 | -0.271 |
| **10610.32** | 0.8411 | 0.0407 | 0.8003 | 0.629 | -0.11 | 0.0606 | 0.8814 | -0.3226 | 0.6165 | 0.7964 | 0.4125 | 0.8367 |
| **11318.98** | -0.1242 | -0.202 | -0.4997 | -0.3011 | -0.4157 | -0.6709 | -0.3823 | -0.2001 | -0.6707 | -0.2013 | 0.1907 | -0.4555 |
| **11938.41** | -0.251 | 0.5563 | -0.0333 | -0.3023 | 0.7193 | -0.4402 | -0.1248 | -0.0869 | 0.021 | -0.1252 | 0.3765 | -0.1376 |
| **12056.34** | -0.0345 | 0.3763 | 0.0642 | -0.0608 | 0.1754 | -0.1953 | -0.0007 | -0.1548 | 0.1154 | 0.1283 | 0.6701 | 0.081 |
| **12292.67** | -0.074 | -0.327 | 0.0822 | -0.2468 | -0.7006 | -0.5776 | -0.0419 | -0.3475 | -0.0965 | 0.0751 | 0.0409 | 0.0874 |
| **12430.29** | -0.0661 | -0.478 | -0.525 | -0.0395 | -0.4264 | -0.7577 | -0.0598 | -0.7079 | -0.7608 | 0.0563 | 0.0311 | -0.4501 |
| **12756.78** | 0.1202 | -0.1138 | 0.2735 | -0.0964 | -0.281 | 0.0427 | -0.0799 | 0.5811 | 0.1772 | 0.0634 | -0.0417 | 0.3585 |
| **13010.77** | -0.1035 | -0.492 | 0.3751 | -0.1972 | -0.6036 | -0.1563 | -0.171 | 0.0904 | 0.1524 | -0.0756 | 0.0273 | 0.3262 |
| **13216.04** | -0.0805 | -0.5126 | 0.3532 | -0.3495 | -0.5959 | 0.1858 | -0.1022 | -0.0139 | 0.3318 | 0.0414 | -0.0864 | 0.572 |
| **13514.37** | -0.1025 | 0.3691 | 0.4335 | -0.3163 | -0.0808 | -0.0919 | -0.0713 | 0.0339 | 0.3663 | -0.0181 | 0.2084 | 0.4672 |
| **13839.87** | -0.078 | 0.3628 | -0.344 | -0.2219 | 0.0264 | -0.871 | -0.074 | -0.1936 | -0.5334 | -0.1912 | 0.2945 | -0.363 |
