## Supplementary material for "A MALDI-MS biotyping-like method to address honey bee health status through computational modelling": Table S2

| Index | File Name | Experimental models (*M. l.* / *P. c. c.* / Control) | Classified | Model Classes (1, 2, 3)* | Mismatch | State of analysis |
|---|---|---|---|---|---|---|
| **1** | C:\Documents and Settings\Administrator\Desktop\modeles HBT 2016\Set of indiv spectra for Validation\Individual Spectra-Ml\0_A17-ML\1\1SLin\fid | ***M. l.*** | true | 1 | 1 | - |
| **2** | C:\Documents and Settings\Administrator\Desktop\modeles HBT 2016\Set of indiv spectra for Validation\Individual Spectra-Ml\0_A18-ML\1\1SLin\fid | ***M. l.*** | true | 3 | 0 | - |
| **3** | C:\Documents and Settings\Administrator\Desktop\modeles HBT 2016\Set of indiv spectra for Validation\Individual Spectra-Ml\0_A19-ML\1\1SLin\fid | ***M. l.*** | true | 3 | 0 | - |
| **4** | C:\Documents and Settings\Administrator\Desktop\modeles HBT 2016\Set of indiv spectra for Validation\Individual Spectra-Ml\0_A20-ML\1\1SLin\fid | ***M. l.*** | true | 3 | 0 | - |
| **5** | C:\Documents and Settings\Administrator\Desktop\modeles HBT 2016\Set of indiv spectra for Validation\Individual Spectra-Ml\0_A21-ML\1\1SLin\fid | ***M. l.*** | true | 3 | 0 | - |
| **6** | C:\Documents and Settings\Administrator\Desktop\modeles HBT 2016\Set of indiv spectra for Validation\Individual Spectra-Ml\0_A22-ML\1\1SLin\fid | ***M. l.*** | true | 3 | 0 | - |
| **7** | C:\Documents and Settings\Administrator\Desktop\modeles HBT 2016\Set of indiv spectra for Validation\Individual Spectra-Ml\0_A23-ML\1\1SLin\fid | ***M. l.*** | true | 2 | 1 | - |
| **8** | C:\Documents and Settings\Administrator\Desktop\modeles HBT 2016\Set of indiv spectra for Validation\Individual Spectra-Ml\0_B1-ML\1\1SLin\fid | ***M. l.*** | true | 3 | 0 | - |
| **9** | C:\Documents and Settings\Administrator\Desktop\modeles HBT 2016\Set of indiv spectra for Validation\Individual Spectra-Ml\0_B2-ML\1\1SLin\fid | ***M. l.*** | true | 3 | 0 | - |
| **10** | C:\Documents and Settings\Administrator\Desktop\modeles HBT 2016\Set of indiv spectra for Validation\Individual Spectra-Ml\0_B3-ML\1\1SLin\fid | ***M. l.*** | true | 1 | 1 | - |
| **11** | C:\Documents and Settings\Administrator\Desktop\modeles HBT 201 | **Control** | true | 1 | 0 | - |
| **12** | C:\Documents and Settings\Administrator\Desktop\modeles HBT 201 | **Control** | true | 1 | 0 | - |
| **13** | C:\Documents and Settings\Administrator\Desktop\modeles HBT 201 | **Control** | true | 1 | 0 | - |
| **14** | C:\Documents and Settings\Administrator\Desktop\modeles HBT 201 | **Control** | true | 1 | 0 | - |
| **15** | C:\Documents and Settings\Administrator\Desktop\modeles HBT 201 | **Control** | true | 1 | 0 | - |
| **16** | C:\Documents and Settings\Administrator\Desktop\modeles HBT 201 | **Control** | true | 1 | 0 | - |
| **17** | C:\Documents and Settings\Administrator\Desktop\modeles HBT 201 | **Control** | true | 1 | 0 | - |
| **18** | C:\Documents and Settings\Administrator\Desktop\modeles HBT 201 | **Control** | true | 1 | 0 | - |
| **19** | C:\Documents and Settings\Administrator\Desktop\modeles HBT 201 | **Control** | true | 2 | 1 | - |
| **20** | C:\Documents and Settings\Administrator\Desktop\modeles HBT 201 | **Control** | true | 1 | 0 | - |
| **21** | C:\Documents and Settings\Administrator\Desktop\modeles HBT 201 | **Control** | true | 1 | 0 | - |
| **22** | C:\Documents and Settings\Administrator\Desktop\modeles HBT 201 | **Control** | true | 3 | 1 | - |
| **23** | C:\Documents and Settings\Administrator\Desktop\modeles HBT 201 | **Control** | - | - | - | Excluded not recalibratable |
| **24** | C:\Documents and Settings\Administrator\Desktop\modeles HBT 201 | **Control** | true | 1 | 0 | - |
| **25** | C:\Documents and Settings\Administrator\Desktop\modeles HBT 201 | **Control** | true | 1 | 0 | - |
| **26** | C:\Documents and Settings\Administrator\Desktop\modeles HBT 201 | **Control** | true | 1 | 0 | - |
| **27** | C:\Documents and Settings\Administrator\Desktop\modeles HBT 201 | **Control** | true | 1 | 0 | - |
| **28** | C:\Documents and Settings\Administrator\Desktop\modeles HBT 201 | **Control** | - | - | - | Excluded noise |
| **29** | C:\Documents and Settings\Administrator\Desktop\modeles HBT 201 | **Control** | true | 3 | 1 | - |
| **30** | C:\Documents and Settings\Administrator\Desktop\modeles HBT 201 | **Control** | true | 1 | 0 | - |
| **31** | C:\Documents and Settings\Administrator\Desktop\modeles HBT 201 | **Control** | true | 2 | 1 | - |
| **32** | C:\Documents and Settings\Administrator\Desktop\modeles HBT 201 | **Control** | true | 1 | 0 | - |
| **33** | C:\Documents and Settings\Administrator\Desktop\modeles HBT 201 | **Control** | true | 2 | 1 | - |
| **34** | C:\Documents and Settings\Administrator\Desktop\modeles HBT 201 | **Control** | - | - | - | Excluded not recalibratable |
| **35** | C:\Documents and Settings\Administrator\Desktop\modeles HBT 201 | **Control** | - | - | - | Excluded not recalibratable |
| **36** | C:\Documents and Settings\Administrator\Desktop\modeles HBT 201 | **Control** | true | 3 | 1 | Excluded not recalibratable |
| **37** | Z:\Projets\HBT-CPT analysis and model generation\Exp july 2016\indi | ***P. c. c.*** | - | - | - | Excluded not recalibratable |
| **38** | Z:\Projets\HBT-CPT analysis and model generation\Exp july 2016\indi | ***P. c. c.*** | true | 2 | 0 | |
| **39** | Z:\Projets\HBT-CPT analysis and model generation\Exp july 2016\indi | ***P. c. c.*** | - | - | - | Excluded not recalibratable |
| **40** | Z:\Projets\HBT-CPT analysis and model generation\Exp july 2016\indi | ***P. c. c.*** | true | 2 | 0 | |
| **41** | Z:\Projets\HBT-CPT analysis and model generation\Exp july 2016\indi | ***P. c. c.*** | - | - | - | Excluded noise |
| **42** | Z:\Projets\HBT-CPT analysis and model generation\Exp july 2016\indi | ***P. c. c.*** | - | - | - | Excluded not recalibratable |
| **43** | Z:\Projets\HBT-CPT analysis and model generation\Exp july 2016\indi | ***P. c. c.*** | - | - | - | Excluded not recalibratable |
| **44** | Z:\Projets\HBT-CPT analysis and model generation\Exp july 2016\indi | ***P. c. c.*** | true | 2 | 0 | |
| **45** | Z:\Projets\HBT-CPT analysis and model generation\Exp july 2016\indi | ***P. c. c.*** | - | - | - | Excluded noise |
| **46** | Z:\Projets\HBT-CPT analysis and model generation\Exp july 2016\indi | ***P. c. c.*** | true | 1 | 1 | |
| **47** | Z:\Projets\HBT-CPT analysis and model generation\Exp july 2016\indi | ***P. c. c.*** | - | - | - | Excluded not recalibratable |
| **48** | Z:\Projets\HBT-CPT analysis and model generation\Exp july 2016\indi | ***P. c. c.*** | - | - | - | Excluded not recalibratable |
| **49** | Z:\Projets\HBT-CPT analysis and model generation\Exp july 2016\indi | ***P. c. c.*** | true | 2 | 0 | |
| **50** | Z:\Projets\HBT-CPT analysis and model generation\Exp july 2016\indi | ***P. c. c.*** | true | 2 | 0 | |
| **51** | Z:\Projets\HBT-CPT analysis and model generation\Exp july 2016\indi | ***P. c. c.*** | - | - | - | Excluded not recalibratable |
| **52** | Z:\Projets\HBT-CPT analysis and model generation\Exp july 2016\indi | ***P. c. c.*** | true | 2 | 0 | |
| **53** | Z:\Projets\HBT-CPT analysis and model generation\Exp july 2016\indi | ***P. c. c.*** | true | 2 | 0 | |
| **54** | Z:\Projets\HBT-CPT analysis and model generation\Exp july 2016\indi | ***P. c. c.*** | - | - | - | Excluded not recalibratable |
| **55** | Z:\Projets\HBT-CPT analysis and model generation\Exp july 2016\indi | ***P. c. c.*** | - | - | - | Excluded not recalibratable |
| **56** | Z:\Projets\HBT-CPT analysis and model generation\Exp july 2016\indi | ***P. c. c.*** | true | 2 | 0 | |
| **57** | Z:\Projets\HBT-CPT analysis and model generation\Exp july 2016\indi | ***P. c. c.*** | - | - | - | Excluded noise |
| **58** | Z:\Projets\HBT-CPT analysis and model generation\Exp july 2016\indi | ***P. c. c.*** | true | 3 | 1 | |
| **59** | Z:\Projets\HBT-CPT analysis and model generation\Exp july 2016\indi | ***P. c. c.*** | true | 3 | 1 | |
| **60** | Z:\Projets\HBT-CPT analysis and model generation\Exp july 2016\indi | ***P. c. c.*** | true | 2 | 0 | |
| **61** | Z:\Projets\HBT-CPT analysis and model generation\Exp july 2016\indi | ***P. c. c.*** | - | - | - | Excluded noise |
| **62** | Z:\Projets\HBT-CPT analysis and model generation\Exp july 2016\indi | ***P. c. c.*** | - | - | - | Excluded not recalibratable |
| **63** | Z:\Projets\HBT-CPT analysis and model generation\Exp july 2016\indi | ***P. c. c.*** | true | 2 | 0 | |
| **64** | Z:\Projets\HBT-CPT analysis and model generation\Exp july 2016\indi | ***P. c. c.*** | true | 2 | 0 | |
| **65** | Z:\Projets\HBT-CPT analysis and model generation\Exp july 2016\indi | ***P. c. c.*** | - | - | - | Excluded not recalibratable |
| **66** | Z:\Projets\HBT-CPT analysis and model generation\Exp july 2016\indi | ***P. c. c.*** | - | - | - | Excluded not recalibratable |
| **67** | Z:\Projets\HBT-CPT analysis and model generation\Exp july 2016\indi | ***P. c. c.*** | - | - | - | Excluded not recalibratable |
| **68** | Z:\Projets\HBT-CPT analysis and model generation\Exp july 2016\indi | ***P. c. c.*** | - | - | - | Excluded noise |
| **69** | Z:\Projets\HBT-CPT analysis and model generation\Exp july 2016\indi | ***P. c. c.*** | true | 2 | 0 | |
| **70** | Z:\Projets\HBT-CPT analysis and model generation\Exp july 2016\indi | ***P. c. c.*** | true | 2 | 0 | |
| **71** | Z:\Projets\HBT-CPT analysis and model generation\Exp july 2016\indi | ***P. c. c.*** | - | - | - | Excluded not recalibratable |
| **72** | Z:\Projets\HBT-CPT analysis and model generation\Exp july 2016\indi | ***P. c. c.*** | true | 3 | 1 | |
| **73** | Z:\Projets\HBT-CPT analysis and model generation\Exp july 2016\indi | ***P. c. c.*** | - | - | - | Excluded noise |
