## Supplementary material for "A MALDI-MS biotyping-like method to address honey bee health status through computational modelling": Figure S1

**Fig. S1.**

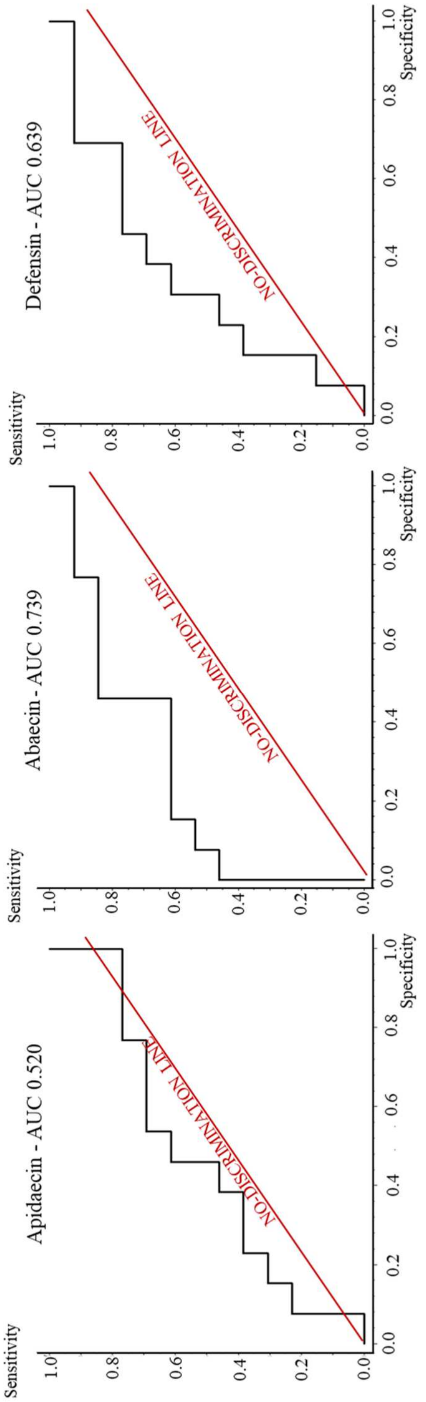


Assessment of ROC curves of Apidaecin, Abaecin and Defensin to discriminate *S. marcescens*- from *S. entomophila*-infected honey bees.
