## Supplementary material for "A MALDI-MS biotyping-like method to address honey bee health status through computational modelling": Figure S2

**Fig. S2.**

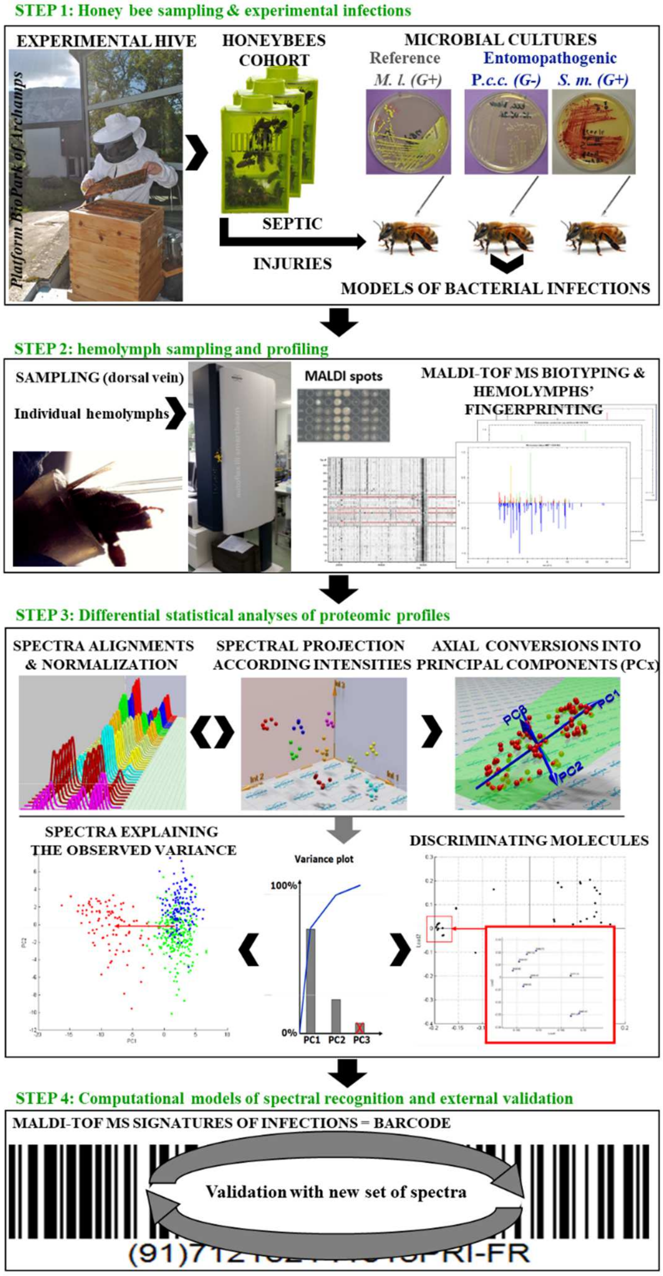


BeeTyping workflow for machine learning data-driven analysis of honey bee infections.

Step 1: Sampling of unchallenged bees (controls) and experimental infection obtained by pricking honey bees with live strains of *P. c. c.* and *M. l.*. Step 2: Individual hemolymph collections followed by MALDI-TOF MS molecular mass fingerprinting, and strain identification by MALDI biotyping. Step 3: Multi-stage processing of MALDI MS fingerprints including recalibration, peak picking, normalization and statistical calculation of individual MS spectra through Principal Component Analysis (PCA) for revealing differential molecular patterns across infection groups. Step 4: Genetic algorithm-based computational model for recognition and classification of honey bee infection using PCA discriminant analysis. Barcodes were built following the molecular fingerprints that discriminate control bees from bees infected either with *M. l.* or with *P. c. c.*.
